# mTORC1-dependent control of Hedgehog signaling mediates nutrient-responsive licensing of intestinal stem cell differentiation

**DOI:** 10.64898/2026.10.29.762847

**Authors:** Gaia Fabris, Simon Biaudet, Atte Hallasaari, Tuuli Kirkinen, Kari Moisio, Krista Kokki, Simon Andersson, Pekka Katajisto, Ville Hietakangas

**Affiliations:** Faculty of Biological and Environmental Sciences, University of Helsinki; Institute of Biotechnology, Helsinki Institute of Life Science, University of Helsinki

## Abstract

How progenitor cells couple their metabolic state to the regulation of fate and differentiation remains a fundamental question in tissue development and regeneration. Here, we identify a nutrient-responsive circuit that links progenitor anabolism to differentiation in the *Drosophila* midgut. We show that the mTORC1 downstream effector PWP1 is necessary and sufficient for differentiation of intestinal stem cell progeny toward the absorptive lineage in fed animals. In progenitors with high mTORC1 activity, PWP1 controls gene expression to promote Hedgehog (Hh) signaling, which cell autonomously drives progenitor differentiation into enterocytes. This circuit is constrained by feedback regulation, as elevated Hh activity inhibits mTORC1 and consequently excludes PWP1 from the nucleus. Disruption of the mTORC1-PWP1-Hh axis causes a progressive failure of enterocyte turnover during aging and reduced lifespan. While the aged midguts retain their barrier function, defective renewal leads to aging-associated deregulation of intestinal metabolism. Collectively, these observations support the concept of growth-dependent licensing of differentiation, whereby the nutrient-responsive anabolic state of a progenitor cell controls its lineage progression, consequently enabling organ renewal.

## INTRODUCTION

A recurring feature of developmental and regenerative systems is that progenitor cell’s growth and metabolic state guide their fate and differentiation (Gong et al., 2023; Meng et al., 2018). This coupling between anabolism and differentiation has been observed in diverse contexts, including neural stem cells, where growth and biosynthetic activity influence the timing of differentiation and lineage progression (Homem et al., 2014), and hematopoietic stem and progenitor cells, in which anabolic and mTORC1-dependent growth programs regulate lineage commitment and differentiation (Kalaitzidis et al., 2012; Signer et al., 2014). Together, these observations support the concept of growth-dependent licensing of differentiation, whereby attainment of an appropriate anabolic state becomes a prerequisite for cell fate commitment. How progenitor cells sense that sufficient anabolic state has been reached and use this information into informing differentiation programs remain, however, poorly understood.

The *Drosophila* midgut provides a powerful model for investigating the coupling between growth and differentiation. The midgut epithelium is organized around three functional cell categories: intestinal stem cells (ISCs), which sustain tissue renewal; transient progenitor cells, comprising enteroblasts (EBs) and enteroendocrine progenitors (pre-EEs); and two terminally differentiated cell types, the nutrient-absorptive enterocytes (ECs) and the hormone-secreting enteroendocrine cells (EEs) (Micchelli & Perrimon, 2006; Ohlstein & Spradling, 2006; Zeng & Hou, 2015). Unlike proliferative ISCs, EBs undergo endoreplication associated with substantial growth before terminal maturation into ECs (Jiang & Edgar, 2011; Xiang et al., 2017). A central regulator of this process is the mechanistic target of rapamycin complex 1 (mTORC1), which integrates nutrient and growth factor signals to promote anabolism and cell growth (Saxton & Sabatini, 2017). In the midgut, mTORC1 is required for nutrient-dependent control of ISC size, which promotes EB fate by activating Delta expression (Mattila et al., 2024). In committed EBs, mTORC1 activity is necessary for growth as well as advancing differentiation towards the EC fate (Kapuria et al., 2012). Both insufficient and excessive mTORC1 activity disrupt intestinal homeostasis, indicating that precise regulation of mTORC1 signaling is essential for productive progenitor differentiation (Amcheslavsky et al., 2011; Haller et al., 2017; Mattila et al., 2024). Despite its established role in EB growth and differentiation, the molecular effectors that couple mTORC1 activity to the signaling pathways that control differentiation remain unclear.

We have previously identified the chromatin-associated protein PWP1 as a nutrient-responsive effector of mTORC1 signaling that promotes growth in *Drosophila* larvae (Liu et al., 2017; Liu et al., 2018). Here we show that PWP1 is necessary and sufficient to promote EB differentiation, downstream of mTORC1 signaling. Through gene regulation, PWP1 controls the activity of the Hedgehog signaling pathway, which cell autonomously promotes EB differentiation into mature ECs. Our data further uncover that high Hedgehog activity in EBs is sufficient to inhibit mTORC1 activity, consistent with a negative feedback mechanism to counterbalance the activity of mTORC1-Hedgehog axis. Disruption of the coupling between EB growth and differentiation compromises intestinal metabolism during aging and reduces organismal lifespan. Collectively, our findings establish a molecular cross-talk enabling growth-dependent licensing of differentiation and underscores its physiological importance in the context of tissue renewal during aging.

## RESULTS

### PWP1 is a nutrient-controlled regulator of intestinal stem cell differentiation

To analyze the role of PWP1 in intestinal stem cell (ISC) regulation, we knocked down its expression in the *Drosophila* adult midgut using the Escargot flip-out (Esg F/O) lineage-tracing system (Jiang et al., 2009), which enables GFP labeling and transgene expression in ISCs and their progeny. PWP1 knockdown (kd) resulted in a significant decrease in the total number of GFP-labeled ISC daughter cells compared to control flies (Figure 1A). Organ-wide computational quantification of the phenotype using the analysis toolkit LAM (Viitanen et al., 2021) showed that ISC activity was specifically impaired in regions R2, R3, and R4 upon PWP1 kd (Figure 1B). Additional MARCM (Mosaic Analysis with a Repressible Cell Marker) analysis – which enables the generation of GFP-positive clones homozygous for a mutation of interest – using a PWP1 null allele revealed a near-complete loss of ISC activity along the midgut (Figure S1A,B). In contrast, the overexpression (oe) of PWP1 in the adult midgut resulted in an increase in ISC progeny number, providing evidence that PWP1 is sufficient to promote ISC activity (Figure 1C,D).

**Figure 1.**
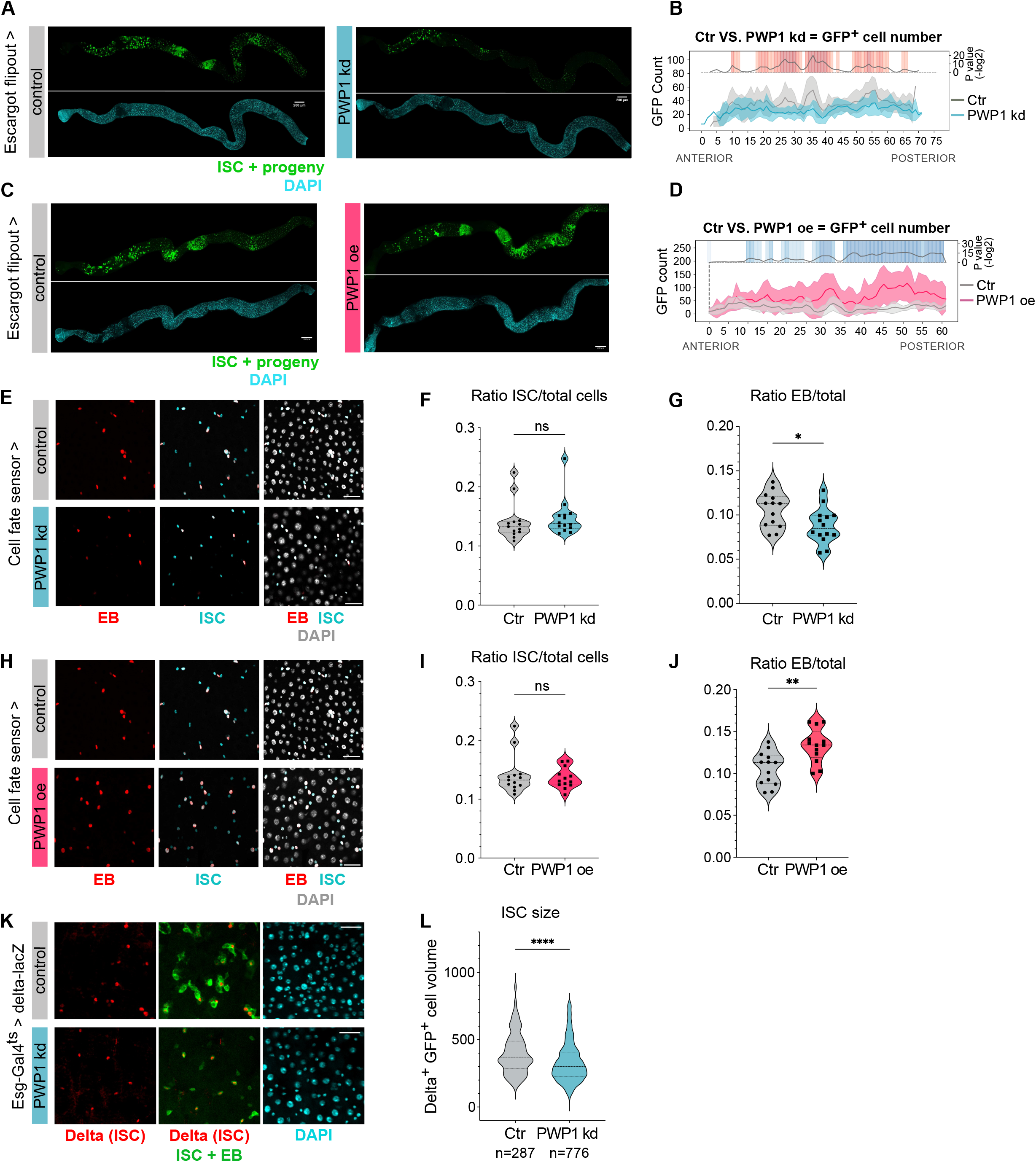
PWP1 is a nutrient-controlled regulator of intestinal stem cell differentiation. **(A)** to **(D)** PWP1 regulates ISC activity. **(A)** Representative images of whole midguts of mated females from Esg-F/Ots > OregonR and Esg-F/Ots > PWP1RNAi, stained with DAPI (cyan) and GFP (green). **(B)** LAM quantification of GFP+ cell number (y axis) along the midgut’s Anterior/Posterior (A/P) axis of control and PWP1RNAi, driven by Esg-F/Ots. Gray and light blue shading correspond to the standard deviation. OregonR n=14, PWP1RNAi n=13. **(C)** Representative images of whole midguts of mated females from Esg-F/Ots > OregonR and Esg-F/Ots > PWP1oe, stained with DAPI (cyan) and GFP (green). **(D)** LAM quantification of GFP+ cell number along the midgut’s A/P axis of control and PWP1oe, driven by Esg-F/Ots. Grey and pink shading correspond to the standard deviation. OregonR n=10, PWP1oe n=9. P values in (B) and (D) were obtained by Mann-Whitney-Wilcoxon U-test with corrections for continuity and multiple testing. Scale bars, 150 μm. (E) to (J) ISC differentiation towards the absorptive lineage is regulated by PWP1. **(E)** Representative images from the posterior midgut of control flies and PWP1RNAi driven by the “cell fate sensor”: grey = all cells; cyan = ISCs; red = EBs. Scale bars, 25 μm. **(F-G)** Quantification of the relative number of ISCs **(F)** and of EBs **(G)** in the posterior midgut. Pooled data from Control n = 16 and PWP1RNAi n = 15 midguts. The data were analyzed using unpaired Welch’s t test, *p = 0.0254, ns indicates no significant difference. **(H)** Representative images from the posterior midgut of control flies and PWP1oe driven by the “cell fate sensor”: grey = all cells; cyan = ISCs; red = EBs. Scale bars, 25 μm. **(I-J)** Quantification of the relative number of ISCs **(I)** and of EBs **(J)** in the posterior midgut. Pooled data from Control n = 16 and PWP1RNAi n = 13 midguts. The data were analyzed using unpaired Welch’s t test, **p = 0.0018, ns indicates no significant difference. **(K-L)** PWP1 regulates ISC size. **(K)** Representative images from the midgut of control flies and PWP1RNAi flies, driven by Esg-Gal4ts. red = Delta+ cells, stained with antibody against beta-galactosidase; GFP = ISCs and EBs; cyan = DAPI. Scale bars, 30 μm. **(L)** Quantification of Delta+-GFP+ cellular volume. Pooled data from OregonR n = 5 and PWP1RNAi n = 7 midguts (the number of cells measured is indicated in the figure panel). The data are analyzed using unpaired Welch’s t test, ****p < 0.0001.

Next, we wanted to address the role of PWP1 in ISC maturation more specifically. The cell fate sensor (Martin et al., 2018), which uses the combinatorial expression of fluorescent reporters to distinguish ISCs from EBs, was combined with Prospero staining to identify enteroendocrine (EE) cells and assess the effects of PWP1 on midgut cell populations (Figure 1E, Figure S1C-F). While PWP1 kd did not affect the relative number of ISCs (Figure 1F) or EE cells (Figure S1C,D), it led to a significant reduction in the relative number of EBs in the posterior midgut (Figure 1G). This finding suggests that PWP1 kd prevents ISCs from differentiating towards the absorptive lineage. Consistent with the loss-of-function data, overexpression of PWP1 increased the proportion of EBs compared to control (Figure 1H-J), while no change in the number of ISCs and EE cells was observed (Figure 1I, Figure S1E,F). The regulation of EB numbers by PWP1 drew our attention to ISC size, as large ISCs were previously shown to be more likely to adopt EB fate (Mattila et al., 2024). Indeed, kd of PWP1 by Esg-Gal4^ts^ (Esg-Gal4; Tub-Gal80) led to a reduction in the nuclear size of Delta^+^ ISCs (Figure 1K,L). Collectively, our data show that PWP1 promotes stem cell differentiation towards the absorptive progenitor EBs.

To study PWP1 expression and regulation in the midgut, we performed immunostainings with the PWP1-specific antibody (Casper et al., 2011). While PWP1 is expressed in nuclei throughout the midgut (Figure 2A), its expression pattern shows strong regional variation. In the posterior midgut (Regions R4-R5), which displays high turnover, PWP1 is primarily expressed in small cells. Co-staining with the stem cell sensor revealed that PWP1 is expressed primarily in ISCs and EBs, and to a lesser extent in EE cells (Figure 2B). As our previous work has shown nutrient regulation of PWP1 expression and its nuclear localization in the larval fat body (Liu et al., 2017), we examined whether feeding status influences PWP1 localization in the midgut. Exposing flies to seven days of nutrient deprivation resulted in clear translocation of PWP1 from the nucleus into the cytoplasm (Figure 2C,D), consistent with starvation-induced inhibition of PWP1’s nuclear function. As PWP1 is a phosphorylation target of mTORC1 (Liu et al., 2017), we tested whether its nuclear localization is mTORC1-dependent, as in fat body cells. Indeed, kd of Raptor, an essential component of mTORC1, led to cytoplasmic translocation of PWP1, similar to that seen with nutrient deprivation (Figure 2E,F). Using the cell fate sensor, we observed that both ISCs and EBs are subject to mTORC1-dependent PWP1 regulation (Figure 2G).

**Figure 2.**
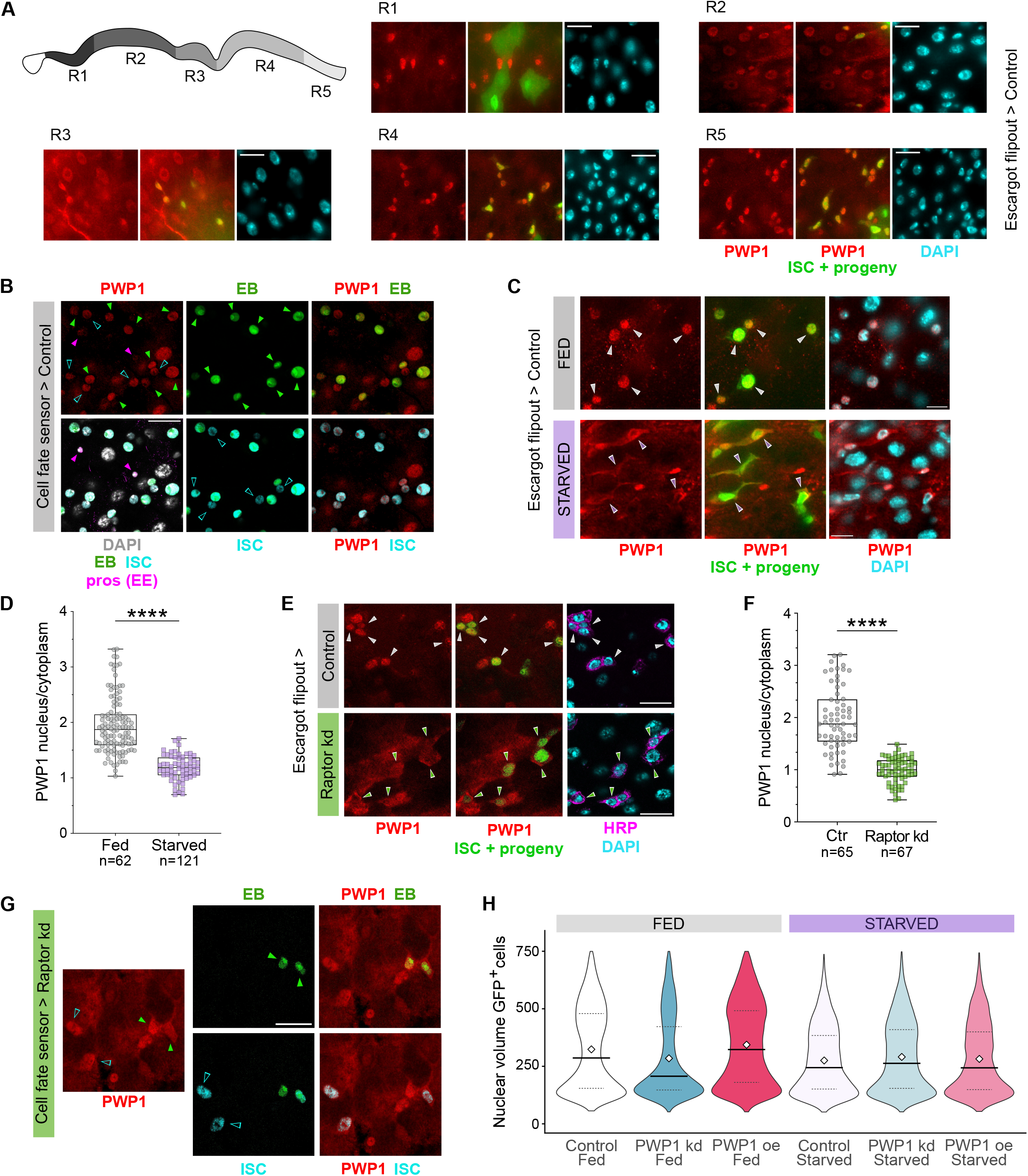
Midgut PWP1 is regulated by mTORC1. **(A)** Representative images of PWP1 expression in the midgut and a schematic view of intestinal regions (R1-5). GFP expression (green) was driven by Esg-F/Ots in wild type (OregonR) female flies. Red = PWP1 antibody staining, cyan = DAPI. Scale bar, 30 μm. **(B)** PWP1 expression across intestinal cell types in the midgut of female control flies, visualized using the “cell fate sensor”: grey = all cells; cyan = ISCs; green = EBs; magenta = EEs (prospero); red = PWP1. Color-matched arrowheads mark representative cells. Scale bar, 30 μm. **(C-D)** PWP1 localization is controlled by nutrients. **(C)** Representative images of PWP1 cellular localization in the midgut of female control flies in fed or starved conditions. GFP expression (green) was driven by Esg-F/Ots, PWP1 expression is marked in red and DAPI in cyan. Color-matched arrowheads mark PWP1 localization in GFP+ cells. Scale bar, 20 μm. **(D)** Quantification of the ratio between PWP1 nuclear and cytoplasmic staining in GFP+ cells. Translocation of PWP1 from the nucleus to the cytoplasm marks its inactivation in the cell. Pooled data from Esg-F/Ots > OregonR Fed n = 3 and Starved n = 3 midguts (the total number of cells measured is indicated in the figure panel). The data are analyzed using unpaired Welch’s t test, ****p < 0.0001. **(E-F)** PWP1 localization is controlled by mTORC1. **(E)** Representative images of PWP1 localization in the posterior midgut of female control (OregonR) flies and RaptorRNAi driven by Esg-F/Ots. GFP (green) = ISCs and their progeny; red = PWP1 antibody staining; magenta = HRP staining (ISCs + EBs); cyan = DAPI. Color-matched arrowheads mark PWP1 localization in GFP+ cells. Scale bar, 30 μm. **(F)** Quantification of the ratio between PWP1 nuclear and cytoplasmic staining in GFP+ cells. Pooled data from OregonR n = 4 and RaptorRNAi n = 5 midguts (the total number of cells measured is indicated in the figure panel). The data are analyzed using unpaired Welch’s t test, ****p < 0.0001. **(G)** PWP1 expression in the midgut of female RaptorRNAi flies, reported by the “cell fate sensor”: cyan = ISCs; green = EBs; red = PWP1. Color-matched arrowheads mark representative cells. Scale bar, 30 μm. **(H)** Nuclear volume distribution of GFP+ cells from control flies (OregonR), PWP1RNAi and PWP1oe driven by Esg-F/Ots, in fed and starved condition. OregonR Fed n = 10, OregonR Starved n = 11, PWP1RNAi Fed n = 9,

Based on the fasting-responsive inhibition of PWP1 nuclear localization, we hypothesized that progenitor cells in nutrient-deprived animals would be unresponsive to changes in PWP1 expression. Nuclear volume of the GFP^+^ cell population was used as a proxy for midgut cell types, with small, mid-size, and large nuclei corresponding to ISCs/EEs, EBs, and ECs, respectively. Consistent with the earlier findings, PWP1 kd increased the proportion of small cells, whereas the relative proportions of mid-size (EBs) and large (ECs) cells decreased compared to controls (Figure 2H, Figure S2A,B). Conversely, PWP1 oe increased the proportion of mid-size cells. However, after seven days of starvation, PWP1 modulation no longer affected the relative proportions of ISCs and their progeny, as the cellular composition remained comparable across all genotypes. Collectively, our findings are consistent with the model that prolonged nutrient deprivation functionally inactivates PWP1, making the progenitors insensitive to the modulation of its expression levels.

### PWP1 directly regulates the Hedgehog signaling pathway

To investigate how the chromatin-associated regulator PWP1 controls ISC differentiation through transcriptional regulation, we utilized the TaDa method (Marshall et al., 2016), employing stem cell specific expression of PWP1 fused with the bacterial Dam methylase (Figure S3A). TaDa analysis identified 827 genes directly bound by PWP1 (Figure S3B, Supplementary Table S1). Over-representation analysis (ORA) data revealed PWP1’s association with genes enriched for innate immune signaling, cytoskeletal and extracellular matrix organization and core developmental signaling cascades (Figure S3C), indicating broad genomic occupancy. In parallel, we performed RNA sequencing of FACS-purified, GFP-labeled ISCs from control flies and flies expressing PWP1-RNAi under the control of ISC-specific driver (Esg-Gal4; Su(H)-Gal80) to identify PWP1-regulated genes prior to the onset of EB differentiation. This uncovered 391 differentially expressed genes (Figure 3A, Figure S3B, Supplementary Table S2) enriched primarily for translation, ribosome biogenesis, nonsense-mediated decay (NMD) and mitochondrial respiration, consistent with previously described roles of PWP1 (Liu et al., 2017). Intriguingly, the Hedgehog-related signal was consistent across analyses, emerging both from the whole-DEG pathway enrichment analysis (Figure S3D,E) and from the downregulated-only network and treeplot (Figure 3B,C), supporting Hedgehog pathway as potential axis modulated by PWP1.

**Figure 3.**
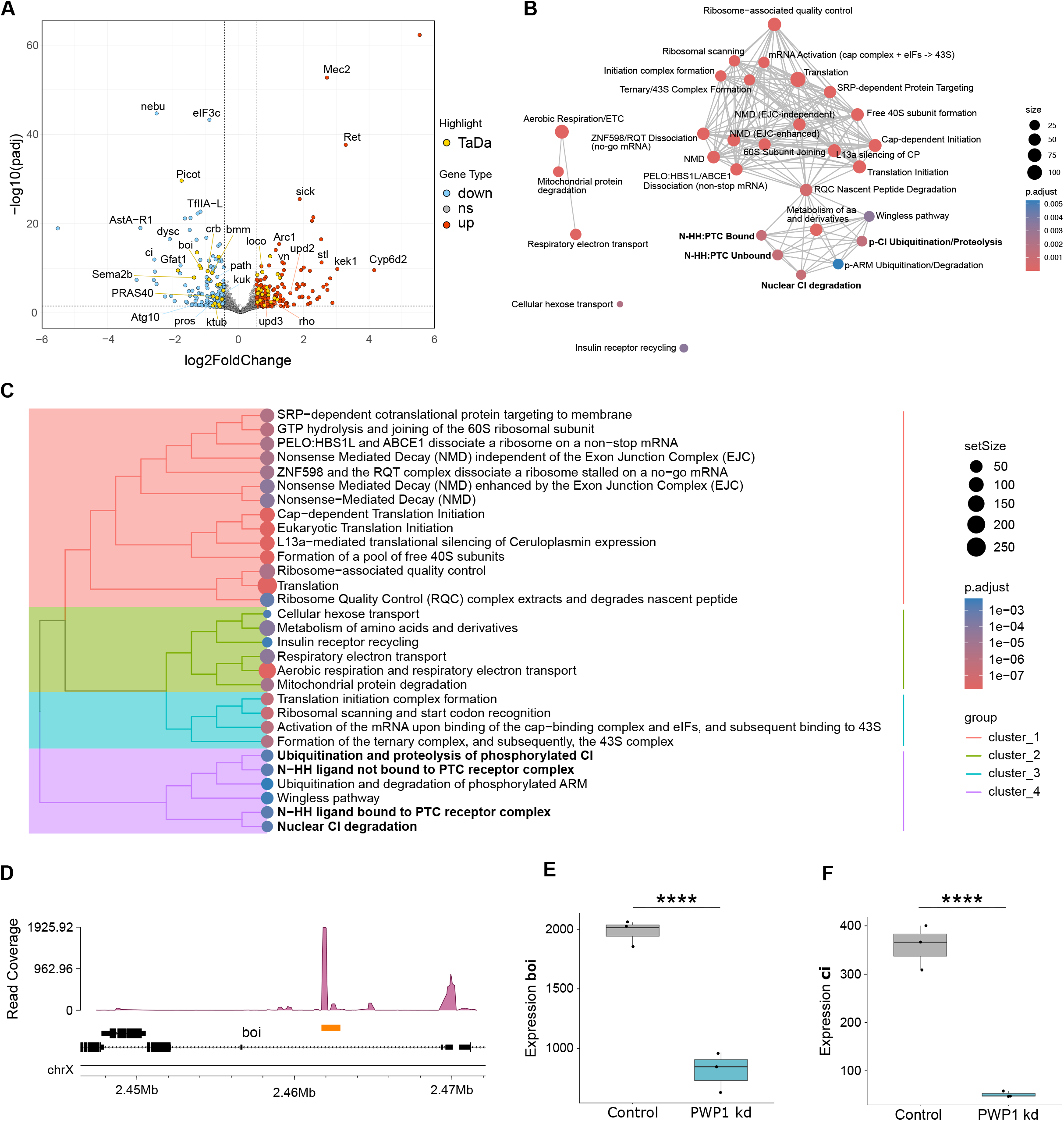
PWP1 regulates components of the Hedgehog signaling pathway. **(A)** Volcano plot of the RNA-seq analysis comparing control (OregonR) and PWP1-kd (PWP1RNAi) flies under the ISC-specific driver (Esg-Gal4; Su(H)-Gal80). Three biological replicates were analyzed for each genotype (OregonR n = 3; PWP1RNAi n = 3), with 80–100 intestines dissected per genotype for each biological replicate. Each dot represents an individual gene. grey = not significant; light blue: down-regulated; red = up-regulated; yellow = DEGs also found from TaDa analysis as PWP1’s direct target. Genes with an adjusted p-value < 0.05 and an absolute fold change > 1 were considered differentially expressed. Genes with very low counts (cpm < 1 in more than one replicate per condition) were filtered out prior to differential expression analysis. **(B-C)** Functional enrichment analysis of downregulated DEGs identified by RNA-seq analysis of PWP1-kd (PWP1RNAi) flies compared with control (OregonR), visualized as a network plot **(B)** and treeplot **(C)**. Hedgehog pathway-related terms are highlighted in bold. Dot size represents the number of genes associated with each enriched pathway and color indicates the adjusted p-value. In the treeplot, related pathways are grouped into clusters according to their similarity. **(D)** TaDa read coverage at the genomic locus of brother of ihog (boi), showing PWP1 binding at the boi locus. The genomic structure and location of the boi gene are shown below the read-coverage profile, highlighted by the orange box. **(E-F)** Normalized RNA-seq expression levels of boi **(E)** and ci **(F)** in control and PWP1-kd (PWP1RNAi) flies. Expression values were normalized using the DESeq2 median-of-ratios method. Each point represents one biological replicate (n = 3 per genotype), and box plots show the distribution of expression values. Statistical significance was assessed using the limma statistical model, with Benjamini–Hochberg correction for multiple testing. ****Adjusted p < 0.0001.

Among the 52 PWP1 transcriptional targets identified in cross-analysis of TaDa and RNAseq datasets (Figure S3B), we identified *brother of ihog* (*boi*), a coreceptor of the Hedgehog receptor Patched (Figure 3D), whose expression was markedly reduced upon PWP1 knockdown (Figure 3E). Notably, RNAseq also revealed decreased expression of *cubitus interruptus* (ci), the transcriptional downstream effector of Hedgehog signaling (Figure 3F), although no significant chromatin binding of PWP1 was detected in its vicinity.

Together, these findings identify Hedgehog signaling as a previously unrecognized downstream target of PWP1 and a candidate mediator of its role in ISC differentiation.

### Hedgehog pathway activity is regulated by mTORC1-PWP1 axis

The findings of PWP1-mediated regulation of Hh pathway components led us to analyze the Hh pathway activity in the midgut epithelium. To directly assess pathway activity *in vivo*, we took advantage of the patched-lacZ (ptz-lacZ) reporter line, sensitive to Hedgehog pathway activity (Struhl et al., 1997). The reporter was active in the progenitor cells in a region-specific manner, with the strongest expression in posterior regions (Figure 4A, Figure S4A). Interestingly, despite the low Hh activity in the progenitors of the anterior intestine, some Hh activity was detected in the EE cells throughout the midgut (Figure 4A, Figure S4A). Overexpression of the Hh pathway activator Smoothened (Smo) by the Esg-F/O system activated the ptz-lacZ reporter (Figure 4B,C), while its knockdown by RNAi led to a marked decrease in lacZ expression (Fig. 4D), validating its function as a Hh pathway sensor in this setting. To more precisely characterize the cell-type specificity, we used the ptc-lacZ reporter in combination with the cell fate sensor, which revealed that the Hh pathway is active in both ISCs and EBs (Figure 4E).

**Figure 4.**
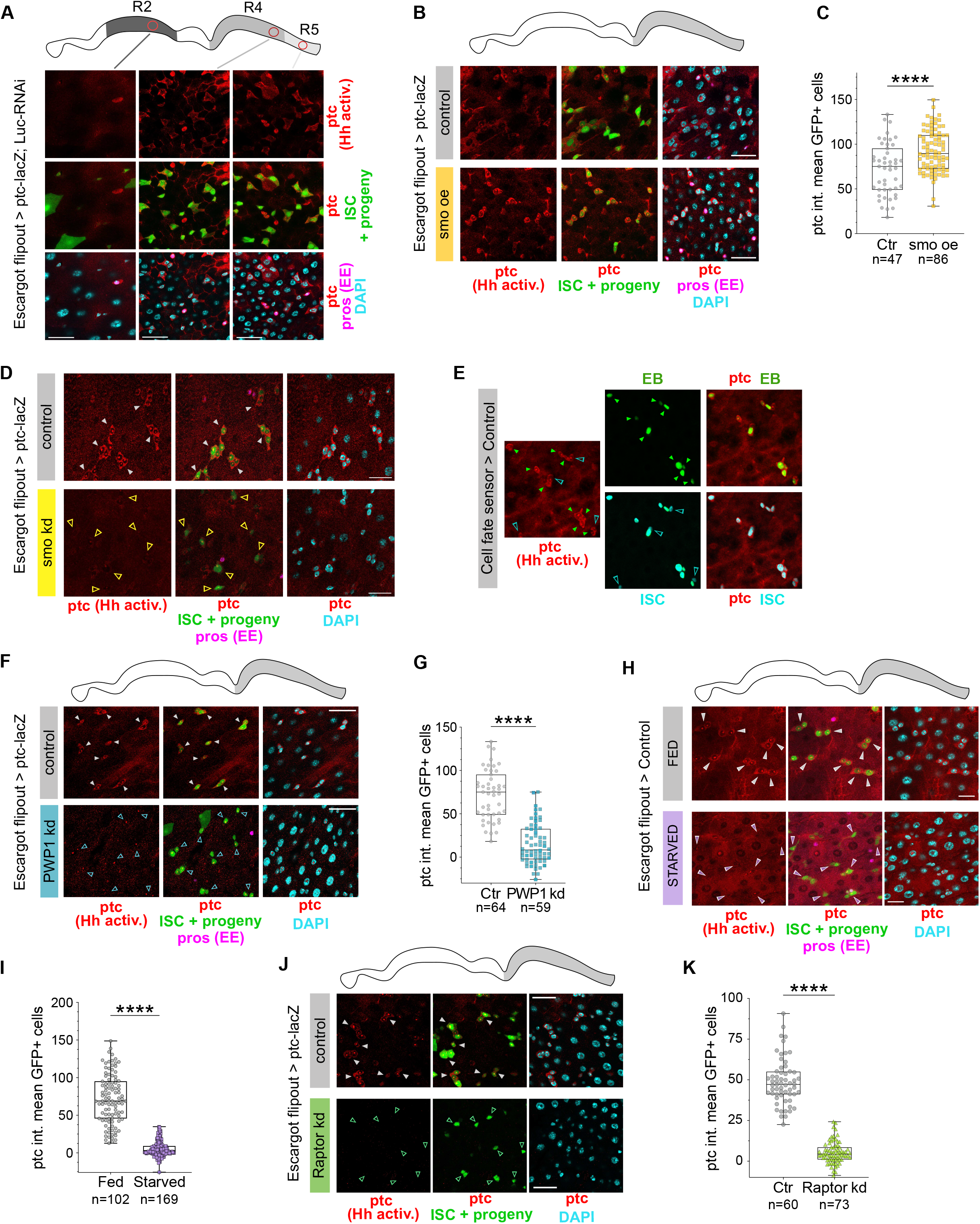
Regulation of the Hedgehog pathway activity by the mTORC1-PWP1 axis. **(A)** Representative images of the Hedgehog (Hh) pathway activation in R2b-c, R4c and R5 of the midgut of control (LucRNAi) flies. patched-lacZ (ptc-lacZ – Hh activation, red) and GFP (ISCs and progeny, green) were driven by Esg-F/Ots (Esg-F/Ots > ptc-lacZ; Luc-RNAi). Prospero markes EEs (magenta) and DAPI marks nuclei (cyan). Scale bar, 25 μm. **(B-C)** smo overexpression (smo oe) activates the Hh pathway. **(B)** Representative images of the posterior midgut of female control (LucRNAi) or smo oe flies, driven by Esg-F/Ots and expressing ptc-lacZ (Esg-F/Ots > ptc-lacZ; Luc-RNAi or UAS-smo). GFP = ISCs and progeny; red = ptc-lacZ activation (beta-galactosidase staining); magenta = EEs (prospero staining); cyan = DAPI. Scale bar, 25 μm. **(C)** Quantification of mean ptc intensity in GFP^+^ cells from control (LucRNAi) and smo oe flies, driven by esg-F/Ots, in the posterior midgut. Pooled data from Luc-RNAi n = 4 and smo oe n = 6 midguts (the total number of cells measured is indicated in the figure panel). The data are analyzed using unpaired Welch’s t test, ****p < 0.0001. **(D)** smo knockdown (smoRNAi) inhibits the Hh pathway. ptc-lacZ and smo-RNAi driven by Esg-F/Ots (Esg-F/Ots > ptc-lacZ; smo-RNAi) was compared to control flies (Esg-F/Ots > ptc-lacZ; Luc-RNAi) in the posterior midgut. GFP = ISCs and progeny; red = ptc-lacZ activation (beta-galactosidase staining); magenta = EEs (prospero staining); cyan = DAPI. Color-matched arrowheads mark representative cells. Scale bar, 25 μm. **(E)** ptc-lacZ expression in the midgut of female control flies, reported by the “cell fate sensor” (cell fate sensor > ptc-lacZ; Luc-RNAi): cyan = ISCs; green = EBs; red = ptc-lacZ activation (beta-galactosidase staining). Color-matched arrowheads mark representative cells. Scale bar, 25 μm. **(F-G)** PWP1 knockdown (PWP1RNAi) inhibits the Hh pathway. **(F)** Representative images of the posterior midgut of female control (LucRNAi) or PWP1RNAi flies, driven by Esg-F/Ots and expressing ptc-lacZ (Esg-F/Ots > ptc-lacZ; Luc-RNAi or PWP1-RNAi). GFP = ISCs and progeny; red = ptc-lacZ activation (beta-galactosidase staining); magenta = EEs (prospero staining); cyan = DAPI. Scale bar, 25 μm. **(G)** Quantification of mean ptc intensity in GFP^+^ cells from control (LucRNAi) and PWP1RNAi flies, driven by esg-F/Ots, in the posterior midgut. Pooled data from Luc-RNAi n = 3 and PWP1RNAi n = 3 midguts (the total number of cells measured is indicated in the figure panel). The data are analyzed using unpaired Welch’s t test, ****p < 0.0001. **(H-I)** Hh pathway activation is controlled by nutrients. **(H)** Representative images of ptc-lacZ expression in the midgut of female control flies (LucRNAi) in fed or starved conditions. GFP expression (green) was driven by Esg-F/Ots, beta-galactosidase staining marks ptc-lacZ activation in red, prospero marks EEs in magenta and DAPI marks nuclei in cyan. Color-matched arrowheads mark ptc-lacZ in GFP+ cells. Scale bar, 25 μm. **(I)** Quantification of mean ptc intensity in GFP^+^ cells from control (LucRNAi) flies in fed and starved condition, under the Esg-F/Ots driver. Pooled data from Fed n = 7 and Starved n = 8 midguts (the total number of cells measured is indicated in the figure panel). The data are analyzed using unpaired Welch’s t test, ****p < 0.0001. **(J-K)** Raptor knockdown (RaptorRNAi) inhibits the Hh pathway. **(J)** Representative images of the posterior midgut of female control (LucRNAi) or RaptorRNAi flies, driven by Esg-F/Ots and expressing ptc-lacZ (Esg-F/Ots > ptc-lacZ; Luc-RNAi or Raptor-RNAi). GFP = ISCs and progeny; red = ptc-lacZ activation (beta-galactosidase staining); cyan = DAPI. Scale bar, 25 μm. **(K)** Quantification of mean ptc intensity in GFP^+^ cells from control (LucRNAi) and RaptorRNAi flies, driven by esg-F/Ots, in the posterior midgut. Pooled data from Luc-RNAi n = 3 and RaptorRNAi n = 3 midguts (the total number of cells measured is indicated in the figure panel). The data are analyzed using unpaired Welch’s t test, ****p < 0.0001.

To test the hypothesis that PWP1 controls Hedgehog pathway activity, we knocked down PWP1 in ISCs and their progeny using Esg F/O, which significantly reduced the expression of the ptc-lacZ reporter (Figure 4F,G). Consistent with the observed nutrient regulation of PWP1, fasting markedly reduced ptc-lacZ activity in progenitor cells (Figure 4H,I). Moreover, inhibition of mTORC1, either through Raptor knockdown or via Rapamycin treatment, resulted in a pronounced decrease in Hh pathway activity in the progenitors of the posterior midgut (Figure 4J,K, Fig. S4B-D). Collectively, these observations showed that the Hh pathway activity in posterior midgut progenitor cells is positively regulated by the mTORC1-PWP1 axis and is responsive to nutrient intake.

### Cell autonomous regulation of absorptive lineage differentiation by Hh signaling

While previous work has shown that Hh signaling in EBs promotes ISC proliferation non-autonomously by upregulating the JAK-STAT ligand Upd2 (Tian et al., 2015), the cell-autonomous role of Hh signaling in midgut progenitors remains poorly understood. We tested this by knocking down the Hh pathway activator Smo or Ptc coreceptor Boi, a direct target of PWP1. Knockdown of either of these Hh pathway components in ISCs and their daughter cells via Esg-F/O resulted in reduced numbers of ISC derived cells in the R4 and R5 midgut regions (Figure 5A-D), mimicking the results obtained with PWP1 knockdown (Figure 2C). Conversely, overexpression of *smo* led to significant increase in ISC progeny (Figure 5E,F). Analysis of their cellular composition in the posterior midgut by the cell fate sensor revealed a strongly reduced numbers of EBs along with moderate reduction in ISCs in smo-kd flies compared to controls (Figure 5G,H, Figure S5A,B).

**Figure 5.**
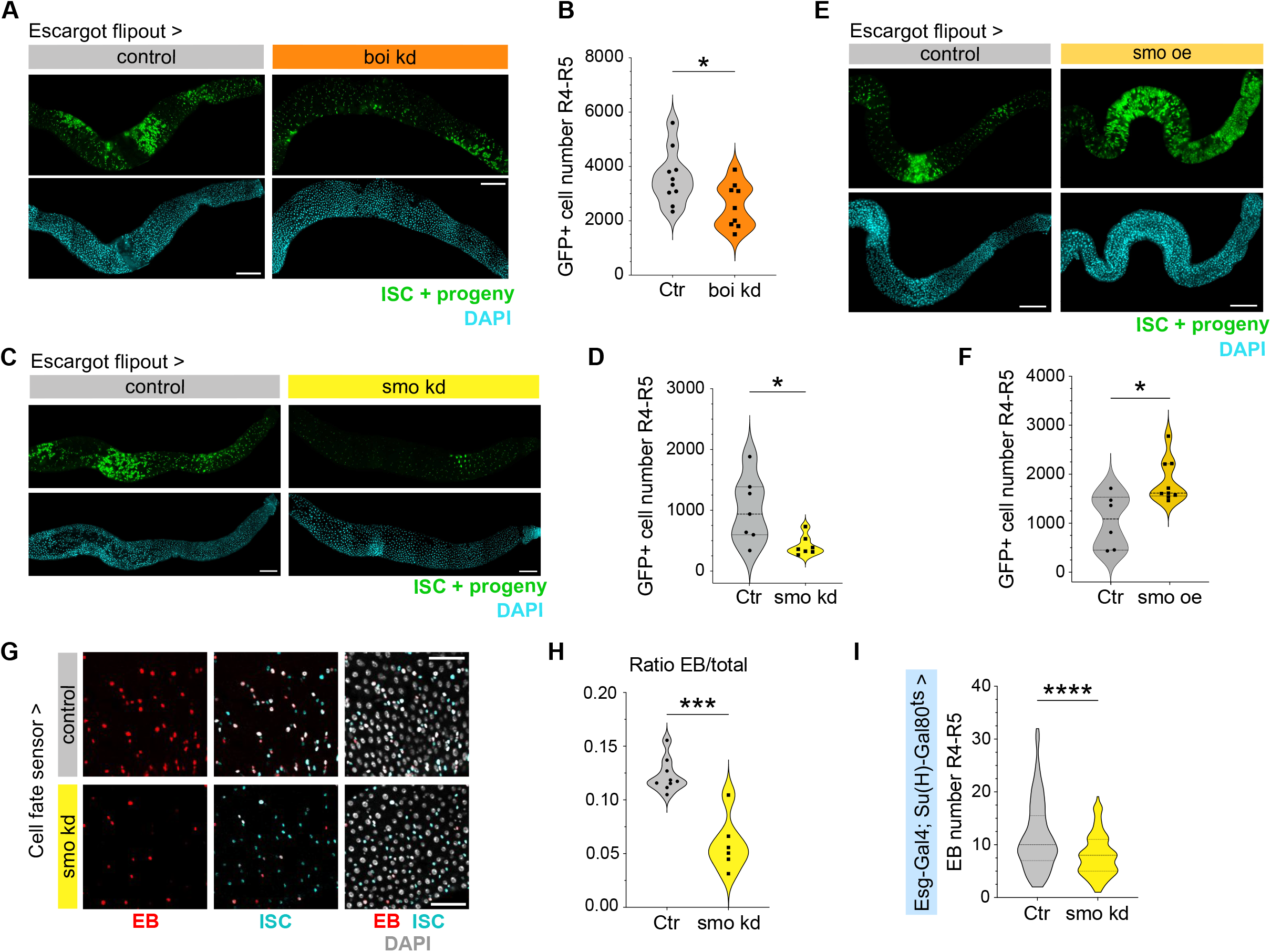
Cell autonomous regulation of the absorptive lineage differentiation by Hh signaling. **(A-B)** boi knockdown (boiRNAi) regulates ISC activity. **(A)** Representative images of the posterior midgut of mated females from control (Esg-F/Ots > LucRNAi) and boi knockdown (Esg-F/Ots > boiRNAi), stained with DAPI (cyan) and GFP (green). Scale bar, 150 μm. **(B)** Quantification of the number of GFP+ cells in the posterior midgut of control vs boi-kd flies under the control of Esg-F/Ots driver. LucRNAi n = 10 and boiRNAi n = 8. The data are analyzed using unpaired Welch’s t test, *p = 0.0247. **(C-D)** smo knockdown (smoRNAi) regulates ISC activity. **(C)** Representative images of the posterior midgut of mated females from control (Esg-F/Ots > LucRNAi) and smo knockdown (Esg-F/Ots > smoRNAi), stained with DAPI (cyan) and GFP (green). Scale bar, 150 μm. **(D)** Quantification of the number of GFP+ cells in the posterior midgut of control vs smo-kd flies under the control of Esg-F/Ots driver. LucRNAi n = 7 and smoRNAi n = 7. The data are analyzed using unpaired Welch’s t test, *p = 0.0270. **(E-F)** smo overexpression (smo oe) regulates ISC activity. **(E)** Representative images of the posterior midgut of mated females from control (Esg-F/Ots > OregonR) and smo overexpression (Esg-F/Ots > smo oe), stained with DAPI (cyan) and GFP (green). Scale bar, 150 μm. **(F)** Quantification of the number of GFP+ cells in the posterior midgut of control vs smo-oe flies under the control of Esg-F/Ots driver. OregonR n = 8 and smooe n = 9. The data are analyzed using unpaired Welch’s t test, *p = 0.0136. **(G-H)** ISC differentiation towards the absorptive cells fate is regulated by the Hedgehog pathway. **(G)** Representative images from the posterior midgut of control flies (OregonR) and smoRNAi driven by the “cell fate sensor”: grey = all cells; cyan = ISCs; red = EBs. Scale bars, 25 μm. **(H)** Quantification of the relative number of EBs in the posterior midgut, from control flies (OregonR) and flies expressing smoRNAi under the control of the “cell fate sensor”. Pooled data from Control n = 9 and smoRNAi n = 6 midguts. The data are analyzed using unpaired Welch’s t test, ***p = 0.0007. **(I)** Quantification of the total number of EBs in the posterior midgut of control flies (OregonR) and smo-kd flies (smoRNAi), under the control of the Esg-Gal4; Su(H)-Gal80ts driver. OregonR n = 27 and smoRNAi n = 27. The data are analyzed using unpaired Welch’s t test, ****p < 0.0001.

We next investigated whether Hh signaling also acts at the ISC stage to regulate their competence to initiate differentiation toward the EC fate. To this end, we knocked down *smo* using the ISC-specific esg-Gal4, Su(H)-Gal80 driver. This impaired ISC differentiation towards the absorptive lineage, as indicated by the reduced number of EBs in Smo-kd flies relative to controls (Figure 5I, Figure S5C). Collectively, these data are consistent with the conclusion that Hh signaling is necessary and sufficient to cell-autonomously promote ISC differentiation towards the absorptive lineage.

### Feedback regulation of mTORC1 by PWP1 and Hh pathway

Given our findings that mTORC1 positively regulates PWP1, and that both are required to sustain Hh pathway activity, we next investigated whether Hh, in turn, feeds back onto this regulatory axis. Indeed, elevated Hh pathway activity through Smo overexpression via Esg-F/O resulted in prominent translocation of PWP1 into the cytoplasm, whereas Smo knockdown did not significantly affect PWP1 nuclear localization (Figure 6A,B). To directly assess the influence of Hh signaling on mTORC1 activity, we performed phospho-4EBP staining (Mattila et al., 2024). Consistent with the effects observed on PWP1, Hh hyperactivation via Smo overexpression led to prominent inhibition of mTORC1 activity, as shown by loss of 4EBP phosphorylation in progenitors, while Smo knockdown did not significantly affect mTORC1 pathway activity compared to controls (Figure 6C,D). Together, these results support a model in which a mTORC1-PWP1 positive feedback loop sustains Hh pathway activity, while Hh hyperactivation, in turn, disrupts this loop by inhibiting PWP1, resulting in decreased mTORC1 activity in progenitors – a negative feedback mechanism that may serve to limit excessive Hh signaling (Figure 6E).

**Figure 6.**
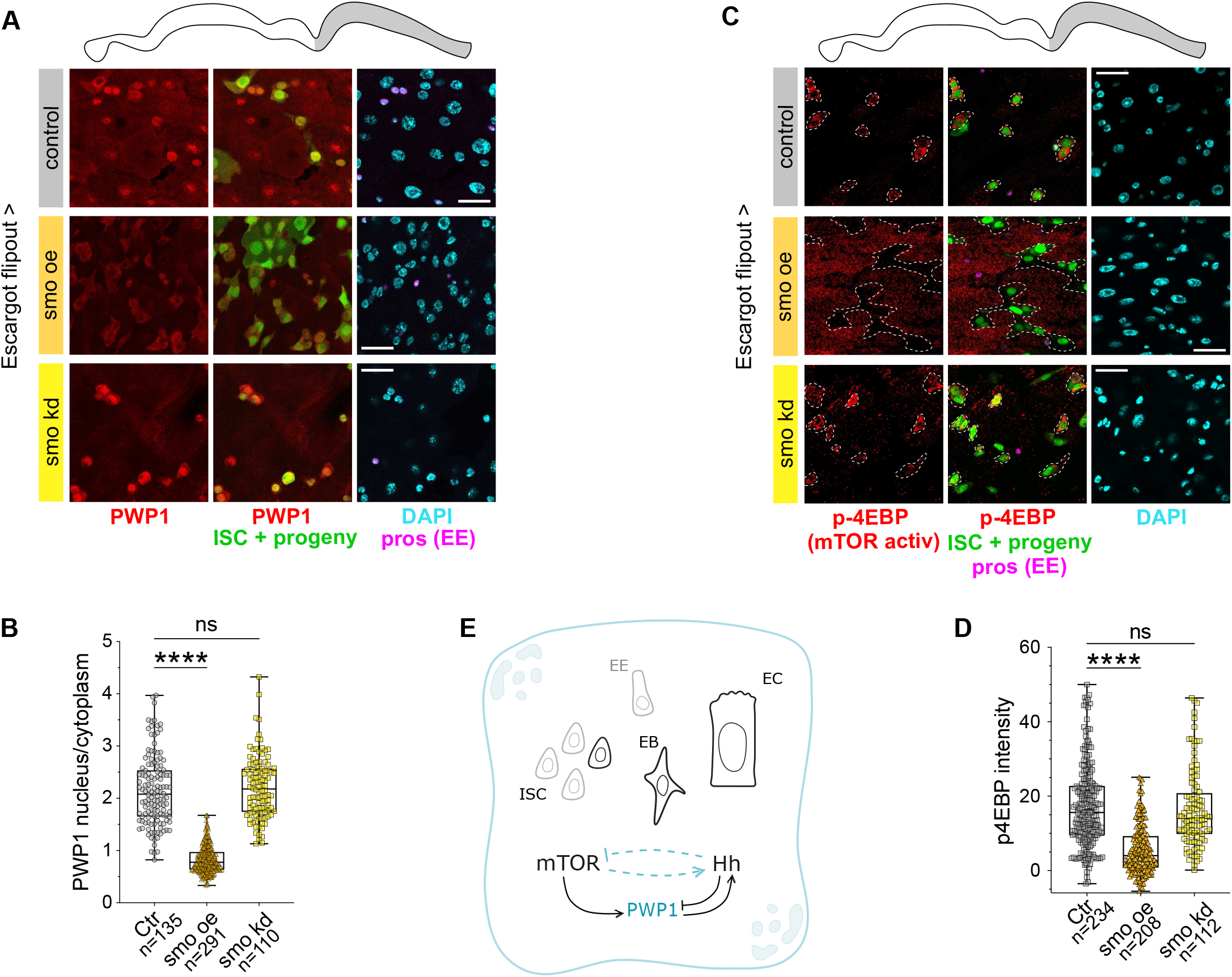
Feedback regulation of mTORC1 by elevated Hh pathway activity. **(A-B)** The Hedgehog pathway regulates PWP1 cellular localization. **(A)** Representative images of PWP1 cellular localization in the posterior midgut of female control flies (OregonR), smo-kd (smoRNAi) and smo-oe (smo oe) driven by Esg-F/Ots. Green (GFP) = ISCs and their progeny; red = PWP1 antibody staining; magenta = EE (prospero staining); cyan = DAPI. Scale bar, 25 μm. **(B)** Quantification of the ratio between PWP1 nuclear and cytoplasmic staining in GFP+ cells. Pooled data from OregonR n = 7, smoRNAi n = 7 and smooe n = 11 midguts (the total number of cells measured is indicated in the figure panel). The data are analyzed using one-way ANOVA followed by Dunnett’s multiple comparisons test, ****p < 0.0001, ns indicates no significant difference. **(C-D)** The Hedgehog pathway regulates mTORC1 activation. **(C)** Representative images of cytoplasmic phospho-4E-BP in the posterior midgut of female control flies (OregonR), smo-kd (smoRNAi) and smo-oe (smo oe) driven by Esg-F/Ots. Green (GFP) = ISCs and their progeny; red = p4E-BP antibody staining; magenta = EE (prospero staining); cyan = DAPI. Dashed lines outline the border of GFP^+^ cells. Cytoplasmic p4E-BP signal is present in control and smo-kd cells, but absent in smo-oe cells. Scale bar, 25 μm. **(D)** Quantification of p4E-BP mean intensity in GFP+ cells. Pooled data from OregonR n = 6, smoRNAi n = 4 and smo oe n = 5 midguts (the total number of cells measured is indicated in the figure panel). The data are analyzed using one-way ANOVA followed by Dunnett’s multiple comparisons test, ****p < 0.0001, ns indicates no significant difference. **(E)** Schematic model of PWP1-mediated regulation of ISC differentiation, highlighting its role as a molecular link between the mTORC1 and Hh pathways.

### PWP1-Hedgehog-mediated turnover sustains midgut metabolism during aging

The Hh pathway has increasingly been associated with tissue regeneration and maintenance during aging, suggesting it may act as a systemic regulator of regenerative and anti-aging processes (Kim et al., 2026). Given the role of the PWP1-Hh axis in regulating ISC differentiation under physiological conditions, we sought to further characterize its relevance in aging flies. As observed in young flies, analysis using the cell fate sensor revealed that PWP1 kd reduced the number of EBs in aged animals (Figure 7A,B), accompanied by a decrease in Hh pathway activity (Figure 7C). When analyzing long-term cumulative turnover in 30-day-old flies using the Esg-F/O lineage tracing system, we observed that chronic (30 days) inhibition of the Hh pathway by Boi kd significantly reduced the number of GFP-positive cells (Figure S7A,B), particularly GFP-positive ECs (Fig. S7C). The effect of long-term PWP1 inhibition on intestinal epithelium turnover was even more dramatic, displaying prominent decrease in GFP-positive cell number compared to control (Figure 7D,E). Most strikingly, the midgut of PWP1 kd flies was almost completely devoid of newly made ECs (Figure 7F). Notably, the nuclear area of 30-day-old GFP-negative ECs was larger compared to control, implying that the lack of EC turnover is compensated by an increase in the size of old cells (Figure S7D).

**Figure 7.**
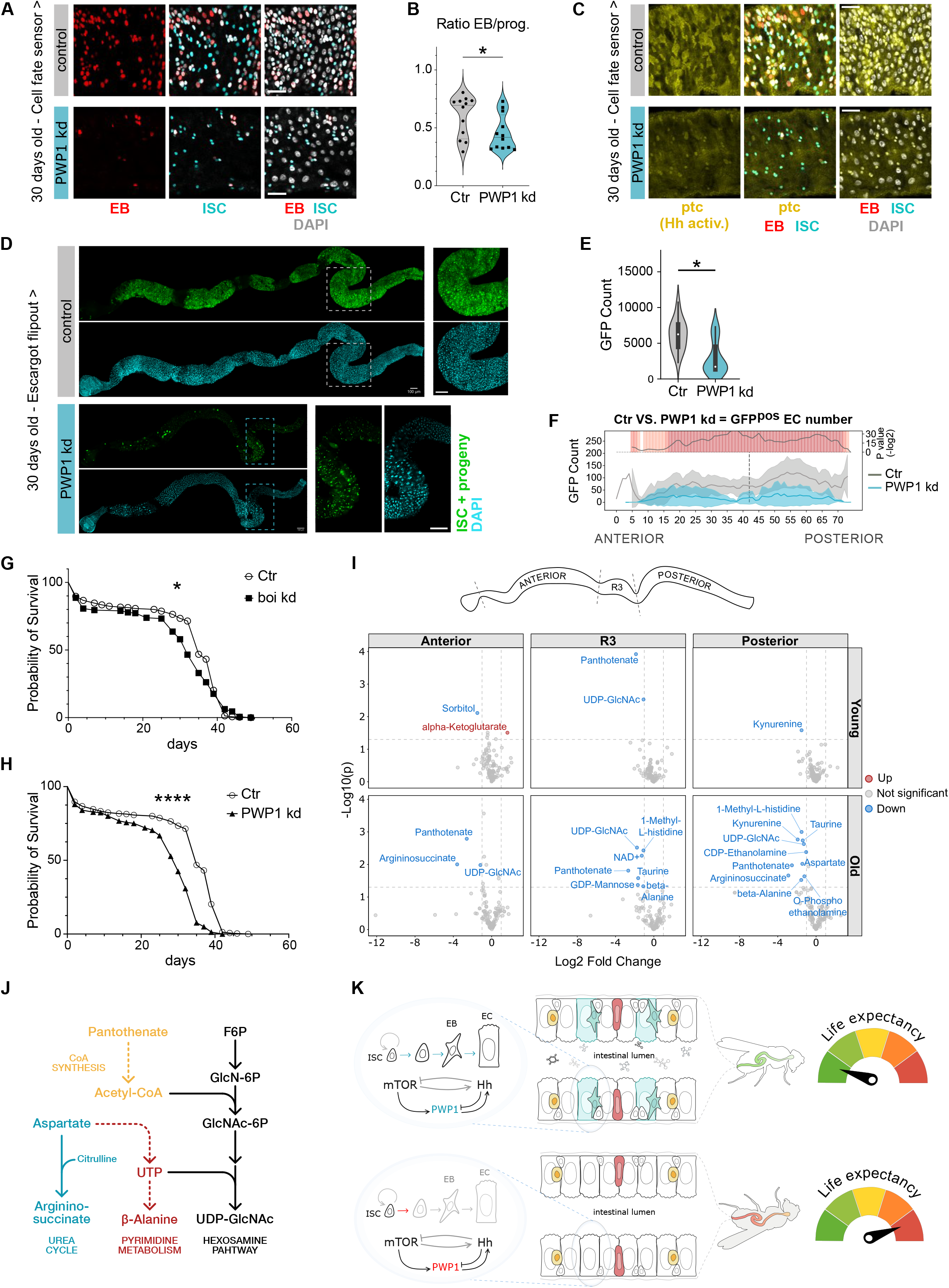
PWP1-Hedgehog axis regulates midgut metabolism and aging. **(A-B)** PWP1 inhibition during aging prevents ISC differentiation towards ECs. **(A)** Representative images from the posterior midgut of control flies (LucRNAi) and PWP1RNAi driven by the “cell fate sensor”: grey = all cells; cyan = ISCs; red = EBs. Scale bars, 25 μm. **(B)** Quantification of the relative number of EBs in 30-days old fly’s posterior midgut. Pooled data from Control n = 12 and PWP1RNAi n = 12 midguts. The data are analyzed using unpaired Welch’s t test, *p = 0.0428. **(C)** PWP1 sustains Hh activity in aged animals. Representative images from the posterior midgut of control flies (LucRNAi) and PWP1RNAi driven by the “cell fate sensor” and expressing ptc-lacZ: grey = all cells; cyan = ISCs; red = EBs, yellow = ptc-lacZ activation (beta-galactosidase staining). Scale bars, 25 μm. **(D)** to **(F)** PWP1 inhibition during aging prevents intestinal turnover. **(D)** Representative images of whole midguts of 30-days old mated females from Esg-F/Ots > OregonR and Esg-F/Ots > PWP1RNAi, stained with DAPI (cyan) and GFP (green). Scale bars whole intestines, 100 μm; scale bars selection, 50 μm. **(E)** Quantification of the number of GFP+ cells. Ctr n = 18 and PWP1RNAi n = 20. ***p < 0.001, P values obtained by Mann-Whitney-Wilcoxon U-test, the thick bar shows the interquartile range and the white dot marks the median. **(F)** LAM quantification of GFP+ EC number along the midgut’s A/P axis of control and PWP1RNAi, driven by Esg-F/Ots. Gray and light blue shading correspond to the standard deviation. OregonR n = 18, PWP1RNAi n = 20. P values were obtained by Mann-Whitney-Wilcoxon U-test with corrections for continuity and multiple testing. **(G-H)** Midgut Hh pathway and PWP1 activity control organismal lifespan. Graphs showing lifespan analysis of control (OregonR) vs boi-kd flies **(G)** or vs PWP1-kd flies **(H)** under the control of Esg-F/Ots driver. (A) OregonR n = 301 and boiRNAi n = 176. **(H)** OregonR n = 301 and PWP1RNAi n = 301. The data are analyzed using the Logrank (Mantel-Cox) test, ****p < 0.0001, *p = 0.0154. **(I)** Volcano plots from regional metabolomic analysis of young and old flies, from control (OregonR) and PWP1-kd (PWP1RNAi) under the control of Esg-F/Ots driver. For each genotype: young n = 3 and old n = 5. The data were analyzed using unpaired Student’s t-test. **(J)** Schematic representation of interconnected metabolic pathways affected in PWP1-kd midguts, highlighting the links between pantothenate/CoA metabolism, aspartate-dependent pyrimidine and urea cycle metabolism, and UDP-GlcNAc biosynthesis. Solid and dashed arrows indicate established metabolic relationships and indirect connections, respectively. **(K)** Schematic illustration of the mTORC1-PWP1-Hh axis in the regulation of ISC differentiation toward the enterocyte (EC) cell fate and its implications for aging. Light blue arrows indicate the progression of ISC differentiation toward the EC fate when PWP1 is active, whereas red text and arrows indicate PWP1 inactivity and impaired ISC differentiation.

Following these findings, we further explored the physiological consequences of impaired PWP1-Hh axis. Consistent with the role of ISC activity in organismal lifespan (Biteau et al., 2010), long-term knockdown of either PWP1 or Boi using the Esg-F/O system resulted in a shorter lifespan compared to control flies (Figure 7G-H). Unexpectedly, however, the intestinal barrier function was not compromised by the impaired turnover. Instead, animals with long-term PWP1 knockdown displayed lower level of intestinal leakage, compared to aged controls, as evidenced by the Smurf assay (Figure S7E).

This finding led us to hypothesize that impaired EC turnover instead manifests through changes in the metabolic function of the midgut. To test this idea, we performed regional metabolomics analysis of the fly intestine in both young and 30-day-old animals. Comparing PWP1 kd to control within each age group, we found a striking asymmetry that mirrors the aging-dependent decline of non-renewed enterocytes: while only a few metabolites were significantly deregulated in young PWP1 kd animals relative to control, old PWP1 kd animals showed roughly a 4-fold increase in downregulated metabolites across all three regions (Figure 7I). This indicates that the metabolic consequences of PWP1 kd are strongly age-dependent, with old animals exhibiting more pronounced metabolic disruption, compared to young animals. Interestingly, the extent of metabolites changes along the midgut showed a pattern that mirrors the regional distribution of Hh pathway activity, which is also highest in R4-R5 (Figure 4A, Figure S4A).

Comparison of regional metabolites changes showed that only pantothenate (vitamin B5) and uridine diphosphate N-acetylglucosamine (UDP-GlcNAc) were reduced across all three regions in old PWP1-kd animals, whereas all other alterations were region-specific (Figure 7I). In young animals, these two metabolites were affected only in R3, indicating that this disruption becomes systemic with age. These deregulated metabolites converge, since pantothenate is a precursor of CoA and acetyl-CoA is used in the biosynthesis of UDP-GlcNAc. UDP-GlcNAc synthesis also relies on pyrimidines, specifically UTP, and concurrent depletion of a pyrimidine precursor, aspartate, and breakdown product, β-alanine, suggest an altered pyrimidine metabolism in old PWP1-kd tissues with aspartate as the constrained input. This is further supported by the depletion of aspartate-derived urea cycle intermediate, argininosuccinate, in the anterior and posterior midgut (Figure 7I). Together, these findings point to impaired pantothenate and aspartate-dependent metabolism as an age-dependent feature of midguts with defective turnover (Figure 7J).

In sum, our data shows that the PWP1-Hh axis is a central regulator of midgut EC turnover, sustaining midgut metabolism rather than epithelial integrity, and that this metabolic regulation supports long-term tissue function and contributes to healthy aging at the organismal level (Figure 7K).

## DISCUSSION

Our findings establish a framework in which the anabolic state of intestinal progenitor cells is coupled to their differentiation, providing a mechanism for growth-dependent licensing of progenitor maturation in the *Drosophila* midgut. We identify mTORC1-PWP1 as an upstream regulator of Hedgehog (Hh) signaling and show that this pathway is required for enteroblast (EB) commitment and maturation into enterocytes (ECs). The circuit is bidirectional: while mTORC1-PWP1 promotes Hh activity, excessive Hh signaling feeds back to suppress mTORC1 activity and PWP1 nuclear localization, thereby constraining the differentiation program. Over the lifespan, this circuit is required for epithelial renewal and maintenance of intestinal metabolic function. Its disruption leads to impaired EC replacement, progressive metabolic deregulation, and shortened lifespan despite preserved epithelial barrier integrity. Together, these findings link nutrient sensing, progenitor growth, differentiation, and tissue aging through a growth-dependent licensing mechanism.

Our previous work identified PWP1 as a nutrient- and mTORC1-responsive chromatin-binding protein required for growth in the *Drosophila* larval fat body (Liu et al., 2017, 2018). PWP1 has also been implicated in cell-fate regulation in other contexts, including differentiation of mouse embryonic stem cells and maintenance of male germline stem cells in *Drosophila* (Casper et al., 2011; Shen et al., 2015). Our findings connect these functions by identifying PWP1 as a molecular link between mTORC1-dependent growth and progenitor cell fate. This extends our previous work showing that mTORC1 regulates ISC size and biases fate toward the absorptive lineage by promoting Delta expression and Notch activation in asymmetric ISC/EB pairs (Mattila et al., 2024). mTORC1 is also highly active in EBs and is required for their maturation into ECs (Kapuria et al., 2012), but how growth signaling is connected to the differentiation machinery was unclear. Our results suggest that mTORC1 acts at two parallel levels of absorptive-lineage development: in ISCs, by promoting Notch-dependent fate specification, and subsequently through PWP1-dependent activation of Hh signaling.

The interaction between mTORC1-PWP1 and Hh is also bidirectional. Although mTORC1-PWP1 is required to sustain Hh activity, excessive Hh activation through Smo overexpression promotes PWP1 cytoplasmic localization and reduces mTORC1 activity, as indicated by decreased 4EBP phosphorylation. The resulting positive cascade coupled to negative feedback may ensure that both anabolic and differentiation signals remain within a productive range. This is consistent with the broader requirement for balanced mTORC1 activity during intestinal homeostasis: excessive mTORC1-driven growth can impair ISC division, while repeated mTORC1 activation during regeneration can contribute to stem-cell depletion and age-related decline (Amcheslavsky et al., 2011; Haller et al., 2017). Together with our previous findings on nutrient-dependent ISC fate regulation (Mattila et al., 2024), these results suggest that the mTORC1-PWP1-Hh circuit coordinates progenitor growth, differentiation, and tissue renewal rather than simply promoting differentiation. How Hh feeds back onto mTORC1 remains an important question for future study.

The identification of Hh as a downstream effector of mTORC1-PWP1 emerged from unbiased analysis of PWP1 targets followed by functional validation. Hh signaling has previously been implicated in midgut regeneration, where tissue damage activates Hh through JNK and Hh promotes ISC proliferation indirectly through Upd2 and JAK-STAT signaling (Tian et al., 2015). Our findings extend this role by showing that Hh is also activated during physiological, nutrient-dependent turnover through mTORC1-PWP1 and is required for intestinal renewal under homeostatic conditions and during aging. This distinction may reflect different inputs converging on Hh. Whereas tissue damage activates Hh through JNK, nutrient availability may engage the pathway through mTORC1-PWP1. Hh could therefore serve as a common downstream effector through which distinct physiological and stress signals adjust intestinal turnover according to tissue demands. Our cell-fate analyses further suggest that Hh acts within the progenitor lineage to promote ISC daughter progression toward the EB state, in addition to its previously described non-autonomous role in stimulating ISC proliferation. The Hh effector Ci is a plausible mediator, as activated Ci can induce Upd2 in EBs (Tian et al., 2015), although the Ci-dependent transcriptional program controlling EB commitment and maturation remains to be defined.

The physiological importance of the mTORC1-PWP1-Hh circuit becomes particularly evident during aging. Chronic depletion of PWP1 or Boi over 30 days caused a near-complete failure to generate new ECs and shortened lifespan. Surprisingly, this profound reduction in epithelial renewal did not compromise barrier integrity as measured by the Smurf assay. Instead, PWP1 depletion suppressed several characteristic age-associated phenotypes, including excessive ISC proliferation and aberrant differentiation, likely because the defective differentiation program prevented the regenerative expansion normally observed in aged controls (Biteau et al., 2008; Choi et al., 2008; López-Otín et al., 2013; Qin et al., 2025). Thus, impaired turnover can preserve aspects of epithelial organization while nevertheless producing severe physiological consequences.

The most prominent consequence of chronic PWP1 depletion was age-dependent metabolic deregulation, particularly in posterior midgut regions where Hh activity and epithelial turnover are high. Depletion of pantothenate and UDP-GlcNAc, together with changes in urea-cycle intermediates, points toward altered aspartate-associated metabolism. Aspartate connects the urea cycle, malate–aspartate shuttle, mitochondrial energy metabolism, and redox homeostasis, and has been implicated in epithelial responses to oxidative stress and in intestinal stem-cell aging (Lokhov et al., 2024; Jin et al., 2025; Luo et al., 2025). Similarly, age-associated decline in pantothenate/CoA metabolism has been linked to ISC dysfunction, while CoA supplementation can reduce age-associated intestinal abnormalities (Liu et al., 2025). UDP-GlcNAc provides an additional potential connection between metabolism and cellular stress. As the end product of the hexosamine biosynthetic pathway, UDP-GlcNAc links nutrient availability to stem-cell behavior and tissue adaptation and serves as the essential substrate for O-GlcNAcylation, a nutrient- and stress-responsive modification involved in signaling, metabolism, aging, and genome maintenance (Mattila et al., 2018; Hart, 2014; Harwood and Hanover, 2014; Na et al., 2020; Liu and Li, 2018). Its depletion in PWP1-depleted aged intestines could therefore affect both metabolic sensing and stress adaptation, although whether impaired UDP-GlcNAc metabolism or O-GlcNAcylation contributes directly to lifespan reduction remains unknown.

Together, our findings identify a molecular circuit that couples mTORC1-dependent growth to Hh-driven differentiation and uses negative feedback to constrain this program. This coupling ensures that progenitor growth, differentiation, and EC replacement remain coordinated with nutrient availability and tissue demands. Disruption of this circuit compromises the coordination between epithelial renewal and metabolic homeostasis, revealing that epithelial renewal is important not only for replacing cells but also for maintaining tissue metabolic competence during aging. Given the conserved roles of mTOR signaling in nutrient sensing, stem-cell regulation, regeneration, and lifespan (Liang et al., 2021; López-Otín et al., 2013; Russell et al., 2011; X. Wei et al., 2019), similar growth–differentiation circuits may represent a broader mechanism linking cellular anabolic state to tissue renewal and metabolic aging.

## MATERIALS & METHODS

### *Drosophila* genetics and husbandry

Flies were maintained at 18°C, on a standard laboratory diet containing agar 0.6% (w/v), malt 6.5% (w/v), semolina 3.2% (w/v), baker’s yeast 1.8% (w/v), nipagin 2.4%, and propionic acid 0.7%. In experiments utilizing the temperature sensitive Gal80^ts^, flies were reared at +18°C and, 2-3 days after hatching, transferred to +29°C for 6 days (young) or for 30 days (old) to inhibit temperature-sensitive Gal80, thereby activating the UAS-GAL4 system in the midgut. Fly stocks used in this study can be found in Supplementary Table S3.

### Dietary treatments

Flies were kept on a chemically defined (holidic) diet (Piper et al., 2013), referred to as fed condition. For starvation, flies were kept on medium containing agar 0.6% (w/v), sucrose 2% (w/v), nipagin 2.4%, and propionic acid 0.7%. The pH of the starvation medium was adjusted to match the holidic diet (pH 6.8) using 5 M NaOH, thus excluding pH as a variable.

### Immunofluorescence

For midgut immunofluorescence staining, intestines were dissected in phosphate-buffered saline (PBS) and fixed in 8% paraformaldehyde for 2 hours. Tissues were washed with 0.1% Triton X-100 in PBS (PBT) and blocked in PBT+1% BSA for 1 hour at room temperature. Following blocking, the tissues were incubated with antibodies against PWP1 (1:500, Atlas HPA038707), Prospero (1:1000, Developmental Studies Hybridoma Bank, MR1A), p4EBP (1:200, Cell Signaling Technology, 2855), β-galactosidase (1:400, MP-Bio 085597-CF), β-galactosidase (1:500, Abcam 9361) or HRP (1:200, Jackson ImmunoResearch Laboratories 2338967) overnight. Secondary antibodies Alexa Fluor™ 568 anti-Rabbit (Invitrogen, A-11011), Alexa Fluor™ 647 anti-Rabbit (Invitrogen, A-21245), Alexa Fluor™ 568 anti-Mouse (Invitrogen, A-11004), Alexa Fluor™ 647 anti-Mouse (Invitrogen, A-21236), Alexa Fluor™ 568 anti-Chicken (Invitrogen, A-11041) were diluted 1:1000 in the blocking buffer and incubated with the tissues at room temperature with mild shaking for 1 hour. Samples were mounted in VECTASHIELD Antifade Mounting Medium with DAPI (Vector Laboratories, H-1200).

### Imaging and image quantification

Whole-mount *Drosophila* midguts, following fixation and immunostaining, were mounted between a microscope slide and an 18 mm coverslip using 0.12 mm spacers. Images of entire intestines were acquired with an Aurox Clarity spinning disc confocal microscope, then processed in ImageJ and segmented using Stardist, as previously described (Viitanen et al., 2021). The resulting raw feature data were exported and imported into R for downstream analysis.

Higher-resolution imaging of selected regions was performed using either a Leica TCS SP8 STED 3X CW 3D microscope or a Leica Stellaris 8 confocal microscope, with manual quantification carried out in ImageJ.

### Targeted Dam ID (TaDa)

Flies were collected 3-4 days post-hatching, and cohorts of 15-20 females and 5-10 males were housed together in new holidic food vials. Flies were then shifted to +29°C for 16 hours to induce transgene expression prior to dissection. For each of five biological replicates, 15 intestines were dissected from female flies expressing either UAS-Dam-PWP1 or UAS-Dam alone under the control of ISC-specific driver (Esg-GAL4; Su(H)-Gal80). The experiment was conducted according to the original protocol (Marshall et al., 2016).

Raw sequencing reads were mapped to the *Drosophila* genome (Dmel_r6.43) and processed with the Marshall lab pipeline (https://owenjm.github.io/damidseq_pipeline/), incorporating some modifications. To reduce the confound of nonspecific methylation typically observed in open chromatin regions, binding profiles of the DAM-fused proteins of interest were normalized against DAM-only DNA binding, with normalization based on kernel density estimation of the log2 GATC ratio. Peaks meeting a stringent FDR threshold (<0.01) were called and assigned to their nearest transcription start site. Peaks overlapping by at least one base pair and detected in 4 or 5 of the 5 UAS-Dam-PWP1 replicates were merged and designated as true binding sites; the resulting pooled peak width thus reflected the combined span of the individual replicate peaks. For visualization purposes, control-subtracted BigWig files were produced using the bigwigCompare tool (deepTools suite (Ramírez et al., 2016), version 3.5.5). These subtracted replicate tracks were then combined into a bedGraph via UCSC’s bigWigMerge, averaged across replicates, and reconverted to BigWig format. The resulting tracks were visualized and compared across conditions using the R package karyoploteR (Gel & Serra, 2017) (version 1.30.0). Genes identified through TaDa were subjected to Over-Representation Analysis (ORA) to identify significantly enriched pathways. ORA was performed using both the clusterProfiler (v4.12.6) and ReactomePA (v1.48.0) R/Bioconductor packages (Yu & He, 2016; Yu, 2026).

### FACS and RNA sequencing

The protocols for FACS and RNA sequencing were adapted from Dutta et al. (2013). Flies were collected 3-4 days post-hatching, and cohorts of 15-20 females and 5-10 males were housed together in fresh starvation media at +29°C to induce transgene expression. After 3 days, flies have been transferred to new holidic food vials and kept at +29°C for 3 additional days prior dissection. For each of three biological replicates, 80 to 100 intestines were dissected from female flies expressing UAS-PWP1-RNAi or from wild type flies (OregonR) under the control of ISC-specific driver (Esg-GAL4; Su(H)-Gal80) in DPBS, and 15 guts at a time were washed and transferred into a pre-coated DNAlow bind tube containing DPBS, placed on ice. Guts were digested with TrypLE™ Select Enzyme (10X) (ThermoFisher Scientific) at 25°C for 30 minutes with shaking at 800rpm. The tissue was dissociated completely by manual agitation, and dissociated cells were centrifuged at 600 x g at 4°C for 10 minutes. Pellet was resuspended in DPBS and filtered with Flowmi® Cell Strainer (70 µm porosity) (Sigma). Propidium iodide (PI) was added to the samples at a final concentration of 2 µg/ml to distinguish nonviable cells (Dutta et al., 2013).

Prior to sorting intestinal stem cells with BD FACSAria™ Fusion flow cytometer, dissociated cells were transferred into Corning™ Falcon™ Test Tube with Cell Strainer Snap Cap (35 µm porosity) (FisherScientific). BD FACSDiva 8.0.2 software was used for sorting. The population of all cells was gated using forward scatter (FSC-A) and side scatter (SSC-A). Viable cells were selected by gating for a PI-negative population, and doublets were removed by gating single cells in dot plots of FSC-W versus FSC-H and SSC-W versus SSC-H. Intestinal stem cells were sorted based on GFP intensity and internal complexity (SSC-A).

RNA was extracted from FACS-sorted ISCs using the Arcturus PicoPure RNA Isolation Kit (Applied Biosystems), following the manufacturer’s instructions, and RNA quality was assessed using an Agilent RNA 6000 Pico chip on the Bioanalyzer.

For mRNA-sequencing, libraries were generated using the TruSeq Stranded mRNA Library Prep Kit (Illumina) and subsequently sequenced with Illumina NextSeq500 technology to an average depth of 20 M reads per sample. The reads were mapped with TopHat to the *D. melanogaster* reference genome (FlyBase R6.10). The differential expression was quantified with DESeq2 (v1.44.0) package (Love et al., 2014) implemented in R/Bioconductor with Benjamini–Hochberg correction for adjusting p values. Genes with an adjusted p-value <0.05 and absolute fold change > 0.5 were considered to be differentially expressed (DE), and they were retained if they had a count greater than 10 in at least two samples in both genotypes. Genes identified as differentially expressed were subjected to Gene Set Enrichment Analysis (GSEA) to identify significantly enriched pathways. GSEA was performed using both clusterProfiler (v.4.12.6) and ReactomePA (v1.48.0) R/bioconductor packages (Yu, 2026; Yu & He, 2016). Pathways with an adjusted p-value < 0.05 with Benjamini–Hochberg correction were considered significantly enriched.

### Metabolomics

Flies were collected 3-4 days after eclosion, and 15-20 females were housed with 5-10 males per vial on holidic medium. To induce transgene expression, flies were shifted to +29°C for either 6 days (young) or 30 days (old) prior to dissection. For each biological replicate, 15 intestines were dissected in DPBS from female flies expressing *UAS-PWP1-RNAi* or from wild-type controls, both under the control of the Esg-F/O driver (3 biological replicates for young flies, 5 for old). Each intestine was manually divided into three regions – anterior (R1-R2), R3, and posterior (R4-R5) – and each region was transferred to a separate 2 ml round-bottom tube containing DPBS on ice. Following collection, samples were centrifuged, the DPBS was removed, and the tissue was snap-frozen in liquid nitrogen and stored at –80°C until processing. On the day of analysis, samples were homogenized using glass beads in a Qiagen TissueLyser (30 s pulses at 30 Hz). Following lysis, ACN:HLO (80:20) extraction buffer was added, samples were briefly vortexed, and the supernatant was analyzed by mass spectrometry.

Samples were analyzed on a Thermo Q Exactive Focus Quadrupole Orbitrap mass spectrometer coupled with a Thermo Dionex UltiMate 3000 HPLC system (Thermo Fisher Scientific). The high-performance liquid chromatography (HPLC) was equipped with a hydrophilic ZIC-pHILIC guard column (20 × 2.1 mm, 5 μm, Merck Sequant). A 5 μL sample was injected into the liquid chromatography-mass spectrometry (LC-MS) instrument after quality controls in randomized order having every tenth sample as blank. A linear solvent gradient was applied in decreasing organic solvent (80%– 35%, 16□min) at 0.15 mL□min–1 flow rate and 45°C column oven temperature. Mobile phases were aqueous 200 mmol per liter ammonium bicarbonate solution (pH∼9.3, adjusted with 25% ammonium hydroxide), 100% acetonitrile, and 100% water. Ammonium bicarbonate solution was kept at 10% throughout the run, resulting in a steady 20 mmol per liter concentration. Metabolites were analyzed using a mass spectrometer with a heated electrospray ionization source using polarity switching and the following settings: resolution of 70,000 at m/z of 200; spray voltages of 3,400 V for positive and 3,000 V for negative mode; sheath gas of 28 arbitrary units (AU) and auxiliary gas of 8∼AU; vaporizer temperature of 280°C; and ion transfer tube temperature of 300°C. The instrument was controlled using Xcalibur 4.1.31.9 software (Thermo Scientific). Metabolite peaks were confirmed using commercial standards (Sigma-Aldrich). Data quality was monitored throughout the run using an in-house quality control cell line extracted similarly to other samples.

Peak integration was done with TraceFinder 4.1 SP2 software (Thermo Scientific). Data processing and analysis were in R version 4.5.1 with tidyverse 2.0.0 package family. Metabolite values were normalized to total metabolite intensity and data from two separate experiments combined into one data set. Metabolites detected reliably in both experiments were analyzed.

### Lifespan

Flies were collected 3-4 days post-hatching and sorted under COL anesthesia. Cohorts of 15-20 females and 5-10 males were housed together in new holidic food vials (10-15 vials per genotype) and maintained horizontally at +29°C for the entire duration of the experiment. Flies were transferred to fresh holidic food every 2-3 days without anesthesia, and female mortality was recorded at each transfer.

### Smurf assay

The Smurf assay was performed in *Drosophila* as previously described (Rera et al., 2011). Flies were collected 3-4 days post-hatching, and cohorts of 15-20 females and 5-10 males were housed together in new holidic food vials (6-10 vials per genotype), maintained horizontally at +29°C for 30 days. Twenty-four hours prior to Smurf scoring, aged flies were transferred onto holidic food supplemented with 2.5% (w/v) Erioglaucine disodium salt. Animals were scored as “Smurf-positive” if the blue dye had diffused throughout the abdomen (early smurf) or the entire body (late smurf) (Martins et al., 2018; Rera et al., 2018).

### Statistical analysis

Statistical analyses were performed in GraphPad Prism 10 (GraphPad Software). For parametric data, unpaired t-test with Welch correction or one-way ANOVA followed by Dunnett’s multiple comparisons test were used. For nonparametric data, the Kruskal-Wallis test or Wilcoxon rank-sum (Mann-Whitney U test) test was used, with multiple testing correction where applicable (FDR < 0.05). For lifespan analysis, the Log-rank test was used. The exact test, sample numbers and P values for each experiment are indicated in the figure legends.

## ACKNOWLEDGMENTS

We thank Heini Lassila, Vid Savnik and Jean Cathala for technical assistance and Onur Deniz for support on computational data management and analysis. Andrea Brand and Catherine Davidson are acknowledged for providing reagents and advice on TaDa. The study was facilitated by the University of Helsinki DNA Sequencing and Genomics Laboratory, Helsinki *Drosophila* core facility (Hi-Fly), and the Light microscopy unit (LMU), with support from Biocenter Finland and Helsinki Institute of Life Science; and by CSC – IT Center for Science, Finland, for providing computational resources.

