## Supplemental figures S1-S7 for "mTORC1-dependent control of Hedgehog signaling mediates nutrient-responsive licensing of intestinal stem cell differentiation"

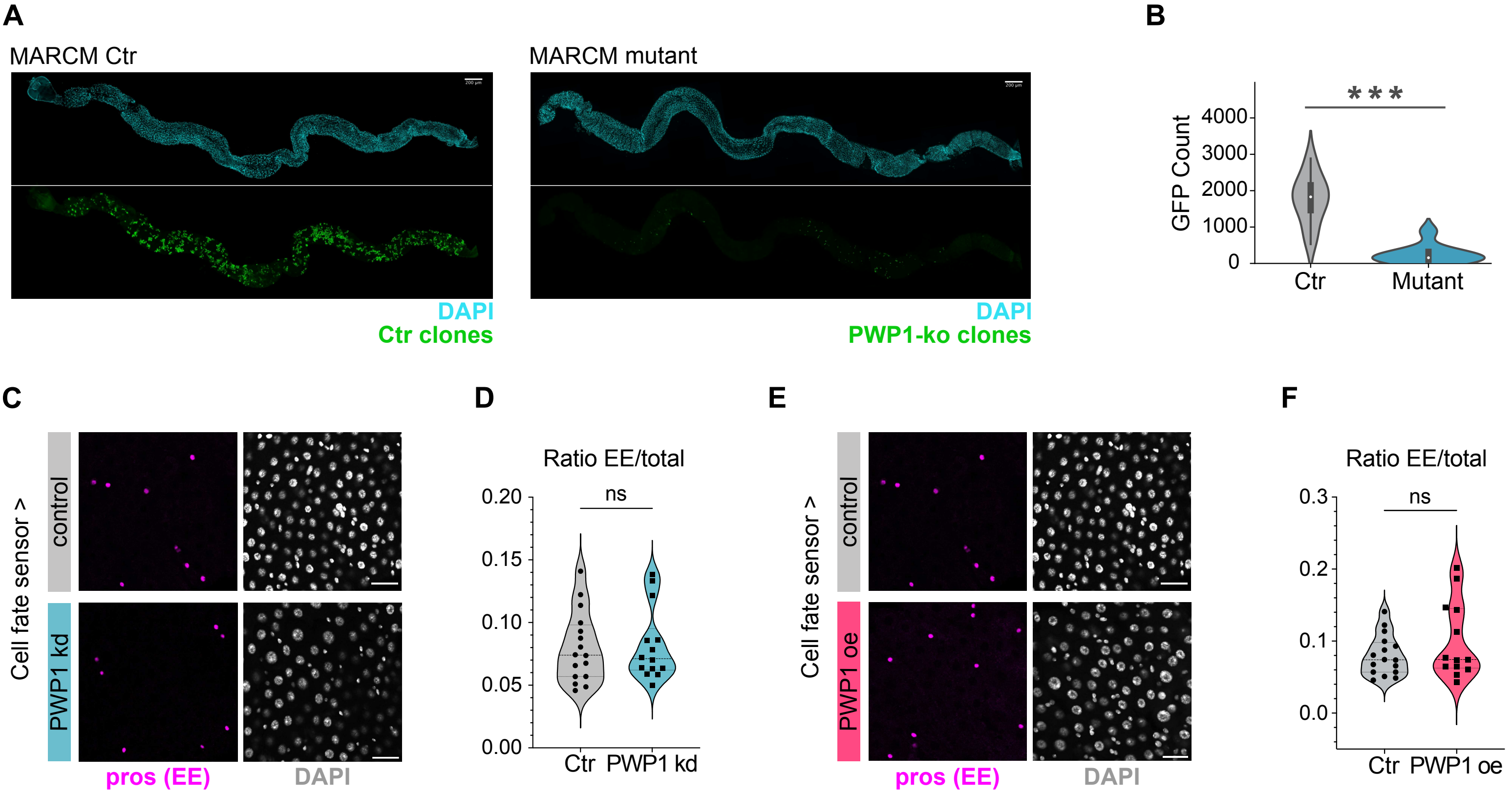


**Supplementary Figure 1.**

**(A-B)** MARCM clones reveal reduced ISC activity in the adult midgut. **(A)** Representative confocal images of GFP-labeled (green) wild-type (Ctr) or PWP1ko mutant MARCM clones in the posterior midgut at 7 days after clone induction (ACI). Flies were heat-shocked at 37°C for 30' at 3–5 days post-eclosion to induce clones. DAPI marks nuclei (cyan). Scale bar, 200 µm. **(B)** Quantification of the number of GFP+ clones. Ctr n = 16 and PWP1ko n = 9. ***p < 0.001, P values obtained by Mann-Whitney-Wilcoxon U-test, the thick bar shows the interquartile range and the white dot marks the median. **(C)** to **(F)** PWP1 modulation does not affect ISC differentiation towards EE cells. **(C)** Representative images from the posterior midgut of control flies and PWP1RNAi driven by the "cell fate sensor": grey = all cells; magenta = Ees (prospero staining). Scale bars, 25 µm. **(D)** Quantification of the relative number of EE in fly's posterior midgut. Pooled data from Control n = 16 and PWP1RNAi n = 15 midguts. The data are analyzed using unpaired Welch's t test, ns indicates no significant difference in the studied parameter. **(E)** Representative images from the posterior midgut of control flies and PWP1oe driven by the "cell fate sensor": grey = all cells; magenta = Ees (prospero staining). Scale bars, 25 µm. **(F)** Quantification of the relative number of EE in fly's posterior midgut. Pooled data from Control n = 16 and PWP1RNAi n = 13 midguts. The data are analyzed using unpaired Welch's t test, ns indicates no significant difference in the studied parameter.

A

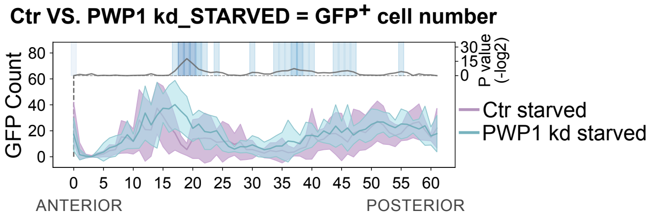


B

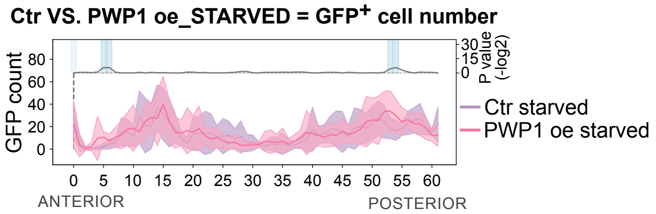


**Supplementary Figure 2.**
**(A-B)** LAM quantification of GFP+ cell number (y axis) along the midgut's Anterior/Posterior (A/P) axis of control and PWP1RNAi **(A)** or control and PWP1oe **(B)** in starved condition, driven by Esg-F/Ots. Lilac, light blue and pink shading correspond to the standard deviation. OregonR Starved n = 11, PWP1RNAi Starved n = 10, PWP1oe Starved n = 10. P values in (A) and (B) were obtained by Mann-Whitney-Wilcoxon U-test with corrections for continuity and multiple testing.

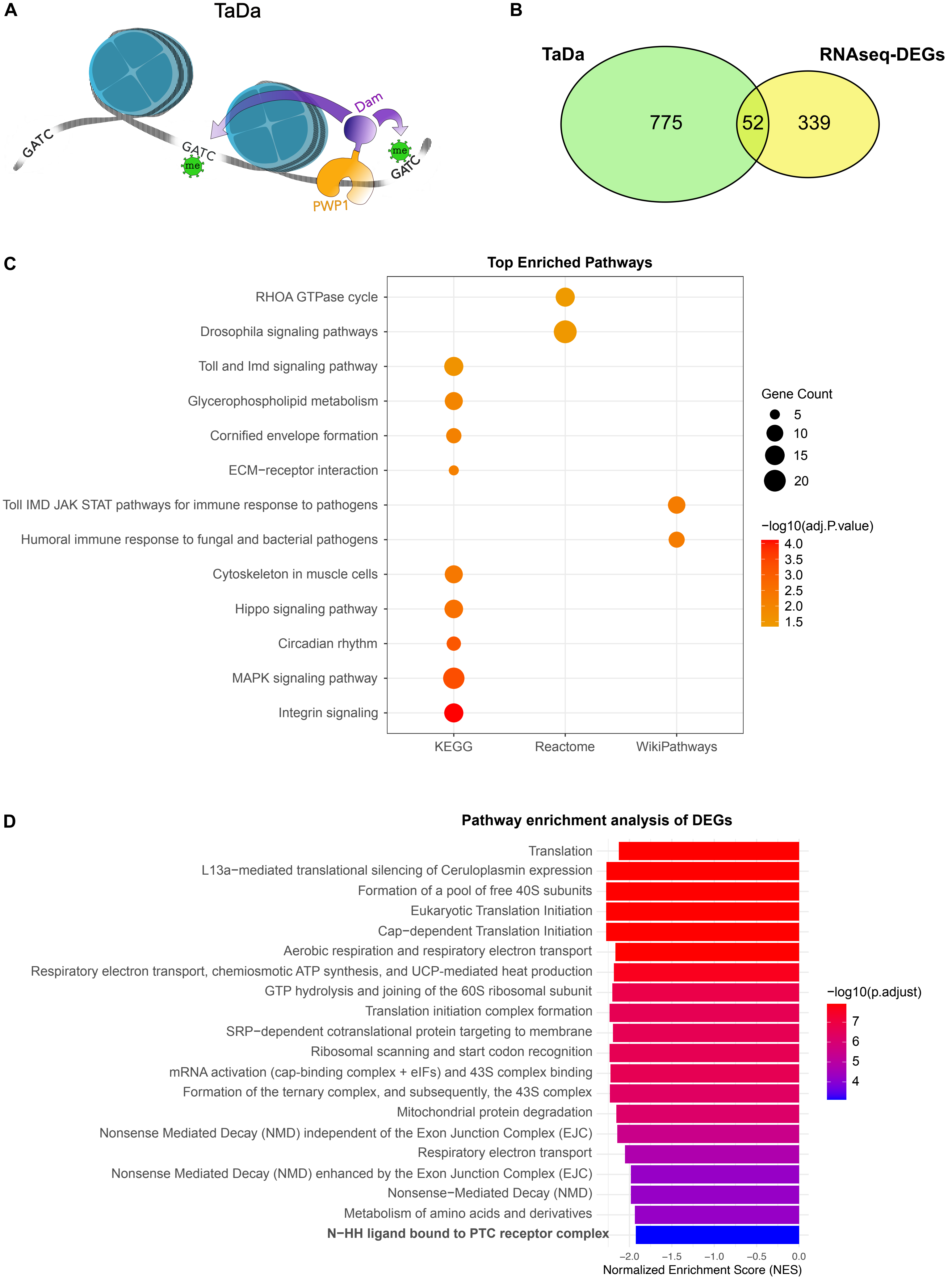
A
TaDa
GATC
GATC
me
Dam
me
GATC
PWP1
B
TaDa
RNAseq-DEGs
775
52
339
C
Top Enriched Pathways
RHOA GTPase cycle
Drosophila signaling pathways
Toll and Imd signaling pathway
Glycerophospholipid metabolism
Cornified envelope formation
ECM−receptor interaction
Toll IMD JAK STAT pathways for immune response to pathogens
Humoral immune response to fungal and bacterial pathogens
Cytoskeleton in muscle cells
Hippo signaling pathway
Circadian rhythm
MAPK signaling pathway
Integrin signaling
KEGG
Reactome
WikiPathways
Gene Count
5
10
15
20
−log10(adj.P.value)
4.0
3.5
3.0
2.5
2.0
1.5
D
Pathway enrichment analysis of DEGs
Translation
L13a−mediated translational silencing of Ceruloplasmin expression
Formation of a pool of free 40S subunits
Eukaryotic Translation Initiation
Cap−dependent Translation Initiation
Aerobic respiration and respiratory electron transport
Respiratory electron transport, chemiosmotic ATP synthesis, and UCP-mediated heat production
GTP hydrolysis and joining of the 60S ribosomal subunit
Translation initiation complex formation
SRP−dependent cotranslational protein targeting to membrane
Ribosomal scanning and start codon recognition
mRNA activation (cap-binding complex + eIFs) and 43S complex binding
Formation of the ternary complex, and subsequently, the 43S complex
Mitochondrial protein degradation
Nonsense Mediated Decay (NMD) independent of the Exon Junction Complex (EJC)
Respiratory electron transport
Nonsense Mediated Decay (NMD) enhanced by the Exon Junction Complex (EJC)
Nonsense−Mediated Decay (NMD)
Metabolism of amino acids and derivatives
N−HH ligand bound to PTC receptor complex
−2.0
−1.5
−1.0
−0.5
0.0
Normalized Enrichment Score (NES)
−log10(p.adjust)
7
6
5
4

**E**

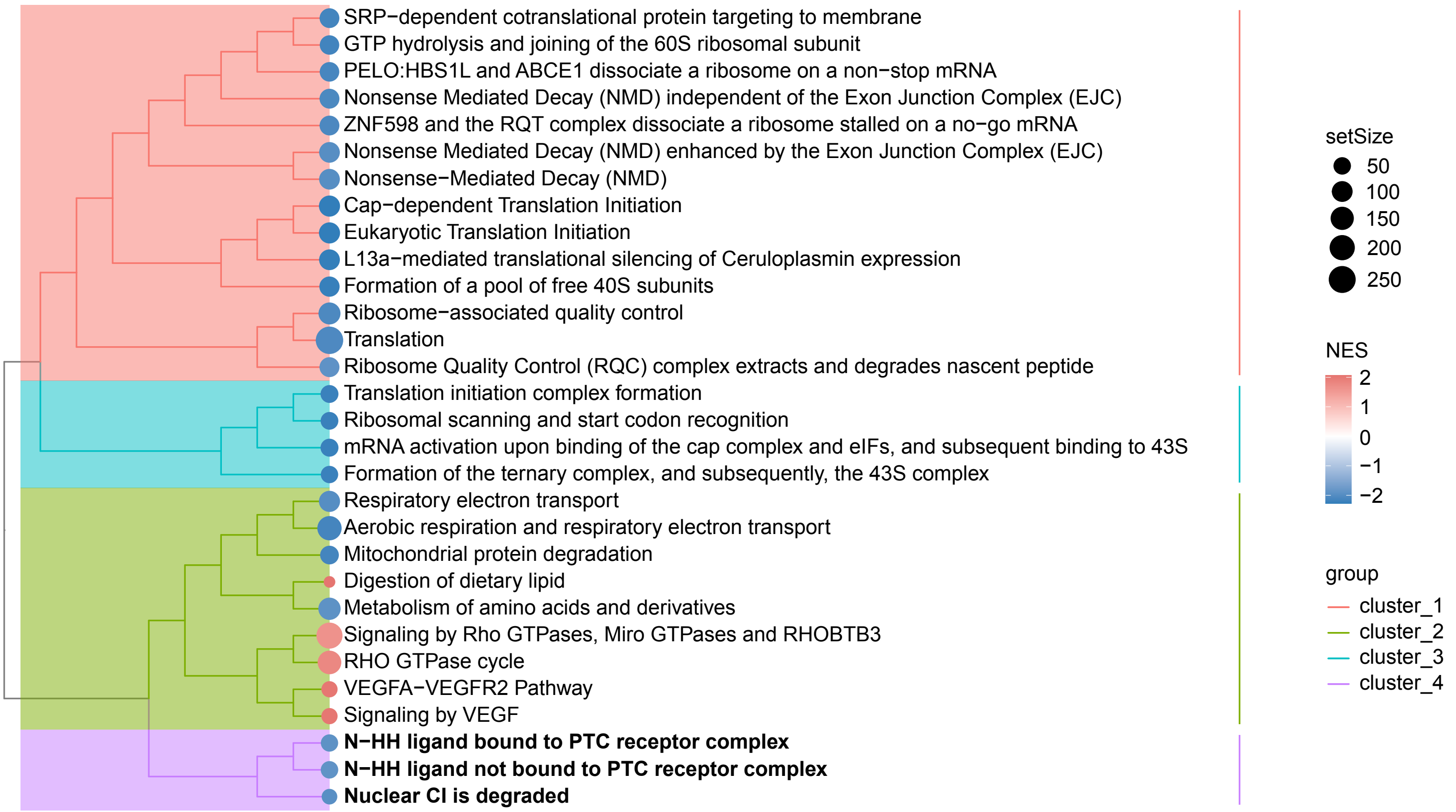


**Supplementary Figure 3.**
**(A)** Schematic of the Targeted DamID (TaDa) approach. PWP1 is fused to a low level of E. coli Dam methylase, which methylates adenines at GATC sites in DNA near PWP1 binding sites. Methylated fragments are then digested, amplified, and sequenced to map PWP1 genome-wide binding profile. **(B)** Venn diagram showing the overlap between genes identified as PWP1 targets by TaDa (green) and differentially expressed genes (DEGs) identified by RNA-seq (yellow). TaDa identified 827 target genes, including 775 TaDa-specific targets and 52 genes overlapping with the RNA-seq dataset. RNA-seq identified 391 DEGs, including 339 RNA-seq-specific genes and 52 genes overlapping with the TaDa dataset. **(C)** Over-representation analysis (ORA) of TaDa target genes, showing the top enriched pathways identified using the KEGG, Reactome, and WikiPathways (WP) databases. **(D–E)** Gene Set Enrichment Analysis (GSEA) of the complete ranked RNA-seq gene list using the Reactome and WikiPathways (WP) databases. Hedgehog pathway-related terms are highlighted in bold. **(D)** Enriched pathways identified by GSEA. **(E)** Treeplot representation of the GSEA results, grouping enriched pathways according to their semantic similarity. Pathways with an adjusted p-value $< 0.05$ were considered significantly enriched. Genes contributing to pathway enrichment are shown according to their direction of expression upon PWP1 knockdown (blue: downregulated; red: upregulated).

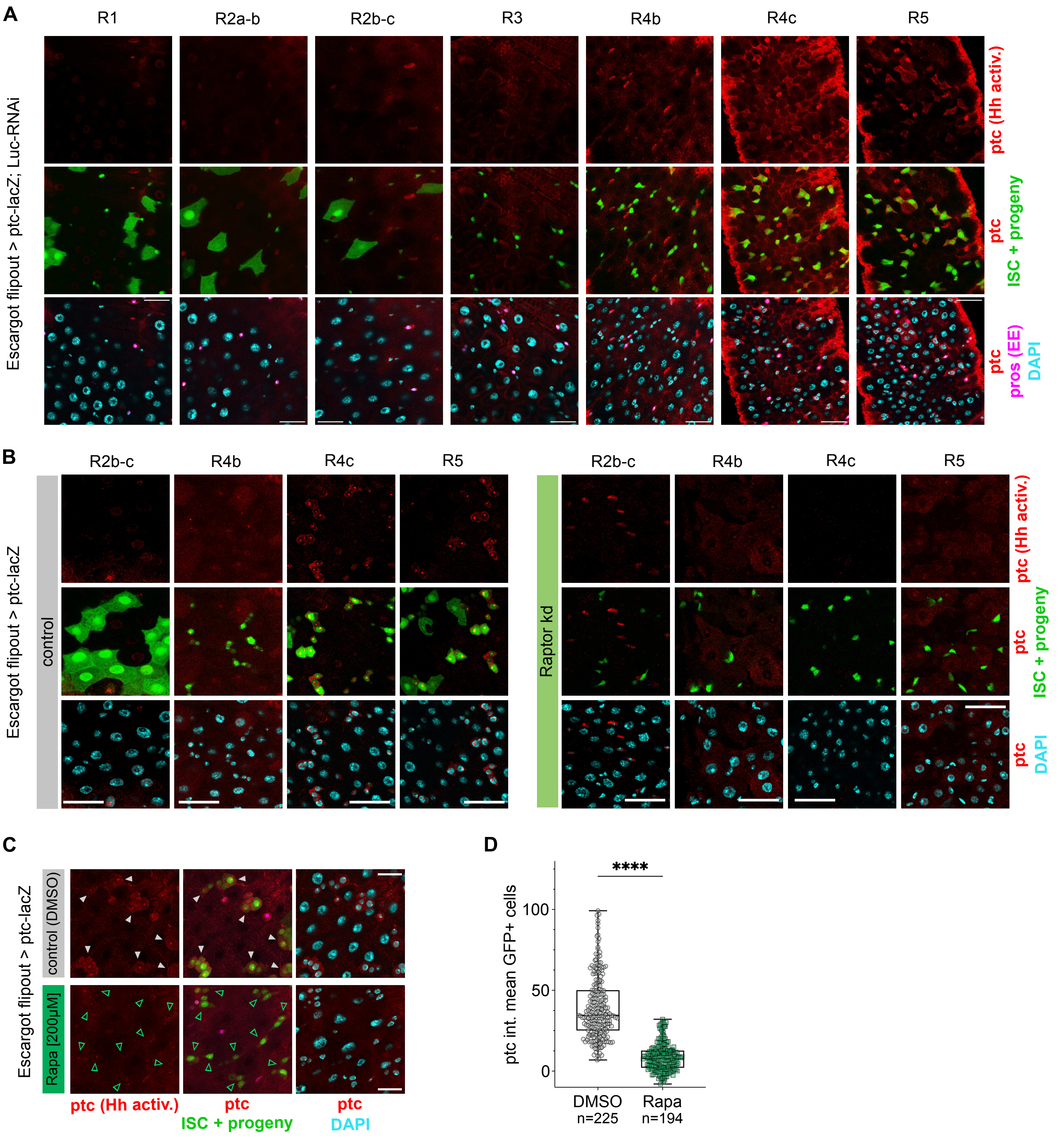


**Supplementary Figure 4.**
**(A)** Representative image of regional differences in Hh pathway activation in female control (LucRNAi) flies, driven by Esg-F/Ots and expressing ptc-lacZ (Esg-F/Ots > ptc-lacZ; Luc-RNAi). GFP = ISCs and progeny; red = ptc-lacZ activation (beta-galactosidase staining); magenta = EEs (prospero staining); cyan = DAPI. Scale bar, 25 μm. **(B)** Representative images of the inhibition of the Hh pathway by mTORC1 in different midgut regions. Posterior midgut of female control (LucRNAi) or RaptorRNAi flies, driven by Esg-F/Ots and expressing ptc-lacZ (Esg-F/Ots > ptc-lacZ; Luc-RNAi or Raptor-RNAi), where green (GFP) = ISCs and progeny; red = ptc-lacZ activation (beta-galactosidase staining); cyan = DAPI. Scale bar, 25 μm. **(C-D)** Rapamycin treatment inhibits the Hh pathway. **(C)** Representative images of the posterior midgut of female control (LucRNAi) flies, driven by Esg-F/Ots and expressing ptc-lacZ (Esg-F/Ots > ptc-lacZ; Luc-RNAi), treated with [200 μM] Rapamycin or DMSO (ctr). GFP = ISCs and progeny; red = ptc-lacZ activation (beta-galactosidase staining); cyan = DAPI. Color-matched arrowheads mark ptc-lacZ in GFP+ cells. Scale bar, 25 μm. **(D)** Quantification of mean ptc intensity in $GFP^+$ cells from control (DMSO) and Rapamycin-treated flies, driven by esg-F/Ots, in the posterior midgut. Pooled data from control (DMSO) n = 4 and Rapa [200 μM] n = 4 midguts (the total number of cells measured is indicated in the figure panel). The data are analyzed using unpaired Welch's t test, ****p < 0.0001.

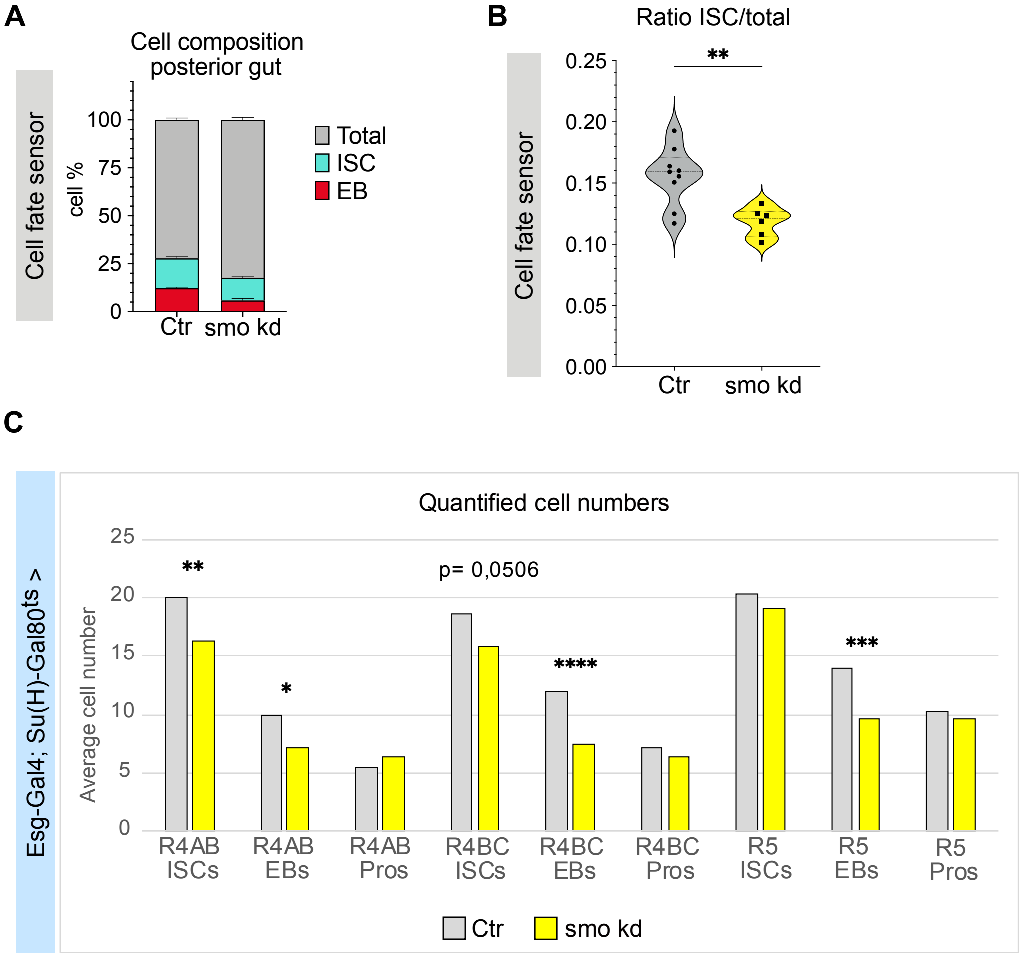


**Supplementary Figure 5.**
**(A-B)** Cell composition **(A)** and quantification of the relative number of ISCs **(B)** in fly's posterior midgut, from control flies (OregonR) and flies expressing smoRNAi under the control of the "cell fate sensor". Pooled data from Control n = 9 and smoRNAi n = 6 midguts. The data are analyzed using unpaired Welch's t test, **p = 0.0014. **(C)** Average cell number per region in control flies (OregonR) and smo-kd flies (smoRNAi), under the control of the Esg-Gal4; Su(H)-Gal80ts driver. OregonR n = 27 and smoRNAi n = 27. The data are analyzed using unpaired t test, **p = 0.0081, *p = 0.0165, ****p < 0.0001, ***p = 0.0029.

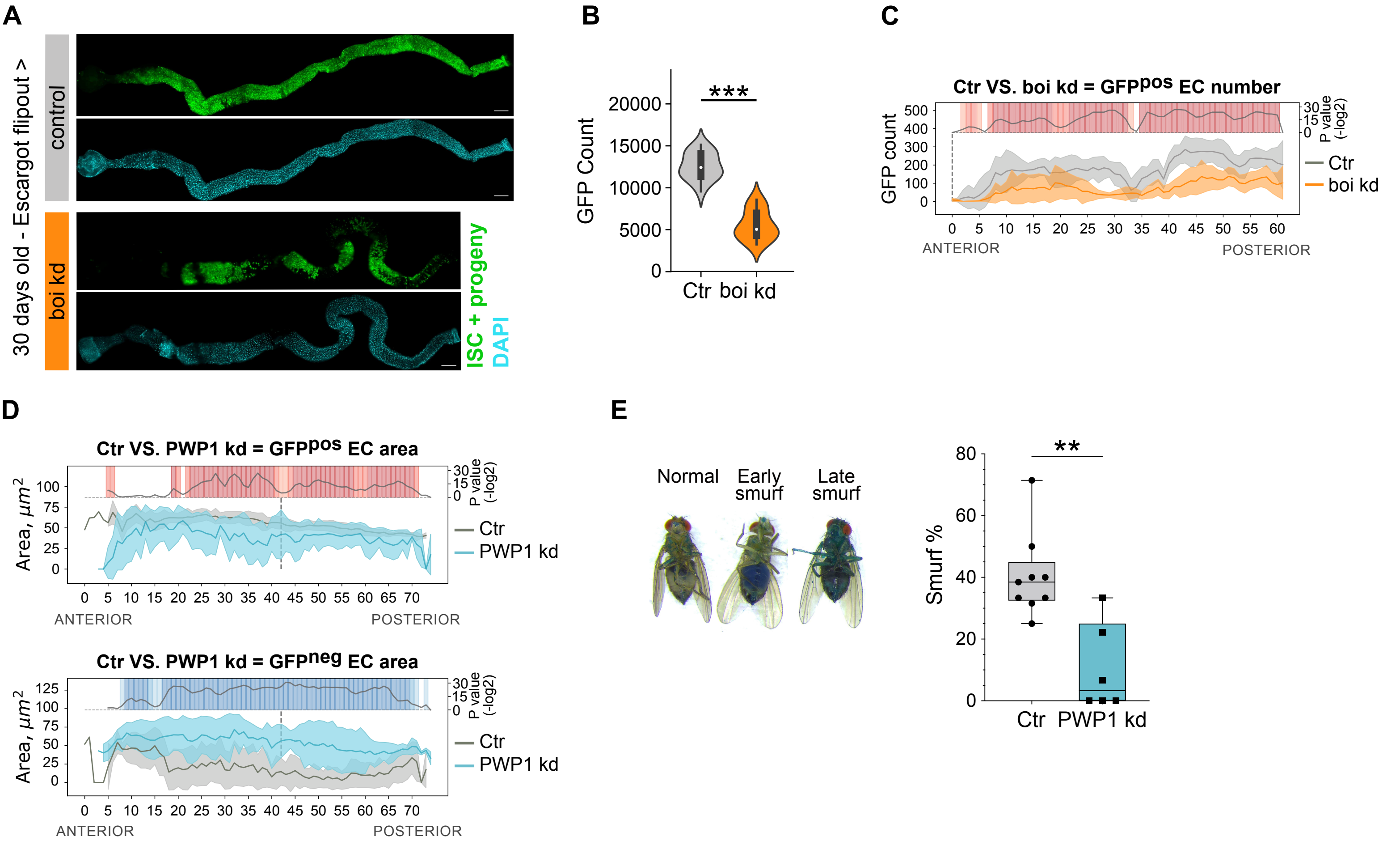


**Supplementary Figure 7.**

**(A)** to **(C)** Inhibition of boi during aging reduces ISC activity. **(A)** Representative images of whole midguts of 30-days old mated females from Esg-F/Ots > LucRNAi and Esg-F/Ots > boiRNAi, stained with DAPI (cyan) and GFP (green). Scale bars whole intestines, 150 µm. **(B)** Quantification of the number of GFP+ cells. Ctr (LucRNAi) n = 15 and boiRNAi n = 13. ***p < 0.001, P values obtained by Mann-Whitney-Wilcoxon U-test, the thick bar shows the interquartile range and the white dot marks the median. **(C)** LAM quantification of GFP+ EC number along the midgut's A/P axis of control (LucRNAi) and boiRNAi, driven by Esg-F/Ots. Gray and orange shading correspond to the standard deviation. LucRNAi n = 15, boiRNAi n = 13. P values were obtained by Mann-Whitney-Wilcoxon U-test with corrections for continuity and multiple testing. **(D)** LAM quantification of GFP+ (top) and GFP- (bottom) EC's nuclear area along the midgut's A/P axis of control (OregonR) and PWP1RNAi, driven by Esg-F/Ots. Gray and light blue shading correspond to the standard deviation. OregonR n = 18, PWP1RNAi n = 20. P values were obtained by Mann-Whitney-Wilcoxon U-test with corrections for continuity and multiple testing. **(E)** Representative image showing the Smurf phenotype and graph of the percentage of Smurfness in 30-days old control (OregonR) vs PWP1-kd (PWP1RNAi) flies, driven by Esg-F/Ots. The data are analyzed using the Mann-Whitney test, **p = 0.0028.
